# Mapping Synthetic Circuit Design Space through Massively Parallel Measurement of Single-Cell Dynamics

**DOI:** 10.64898/2026.10.11.758309

**Authors:** Noah Olsman, Jacob Quinn Shenker, Divya Choudhary, Johan Paulsson

## Abstract

Engineering predictable and robust gene-regulatory networks requires a quantitative understanding of how their dynamics depend on genetic design and environmental conditions. To investigate these relationships, we introduce BARRACUDA, a high-throughput platform that combines combinatorial assembly and programmable perturbations with lineage-resolved time-lapse imaging and optical pooled screening. Experiments using this platform follow approximately one million cell lineages, collectively sampling tens of thousands of circuit designs. We used BARRACUDA to investigate how the robustness of circuit behavior depends on molecular components and their interactions, combining quantitative dynamic measurements with precise genetic and environmental control. Antithetic integral feedback provided a model system linking a defined molecular architecture to theoretical predictions of robust adaptation. Through a hierarchy of increasingly complex libraries, we characterized individual components and used those measurements to model and interpret the dynamics of complete feedback circuits. Circuits built from wild-type inhibitory proteins rejected disturbances but also exhibited sustained overadaptation and fast disturbance-induced oscillations unexplained by a reduced control model. These discrepancies motivated a screen of de novo binders that retained inhibitory function while reducing host-stress response. Selected de novo binders supported strong feedback regulation and robust performance in the rebuilt circuits. Across more than 20,000 recovered feedback-circuit variants, binder identity and controller expression organized regimes of strong regulation and sustained oscillation. Counterintuitively, disturbances sometimes improved compensation rather than compromising it, consistent with nonlinear changes in feedback engagement. Together, these results show how structured libraries and massively parallel measurements of single-cell dynamics can test mechanistic theories across scales of circuit complexity, reveal missing biological interactions, and map the design space of gene-regulatory networks.

## 1 Introduction

A foundational ambition of synthetic biology is to construct regulatory systems whose complexity and capabilities approach those found in nature, while retaining predictable behavior ^1–3^. Landmark synthetic circuits demonstrated how positive feedback can generate bistable switching ^4^, negative feedback can suppress fluctuations in gene expression ^5^, and delayed negative feedback can produce sustained oscillations ^6^. Realizing these behaviors, however, depends on the strengths and timescales of the underlying interactions, not just the connections between components. Even without changing its architecture, a circuit can move between stable regulation and oscillation as expression levels or interaction rates change ^7^. Component activities can also change with molecular context ^8–10^, and new interactions or feedback can expose behavior absent from simpler systems ^11–13^. Building circuits that function robustly beyond a narrow experimental setting therefore requires a quantitative understanding of how their behavior depends on component properties, circuit composition, and environmental inputs.

Mechanistic studies and high-throughput screens offer complementary routes toward this goal. Targeted modeling and experimentation have enabled predictable tuning of bistable switches ^14^ and improved oscillator robustness ^15^, but typically examine limited regions of design space (Fig. 1a, left). Screening explores many more variants and enables the identification of circuits with favorable dynamic properties ^16,17^ (Fig. 1a, center).

**Figure 1.**
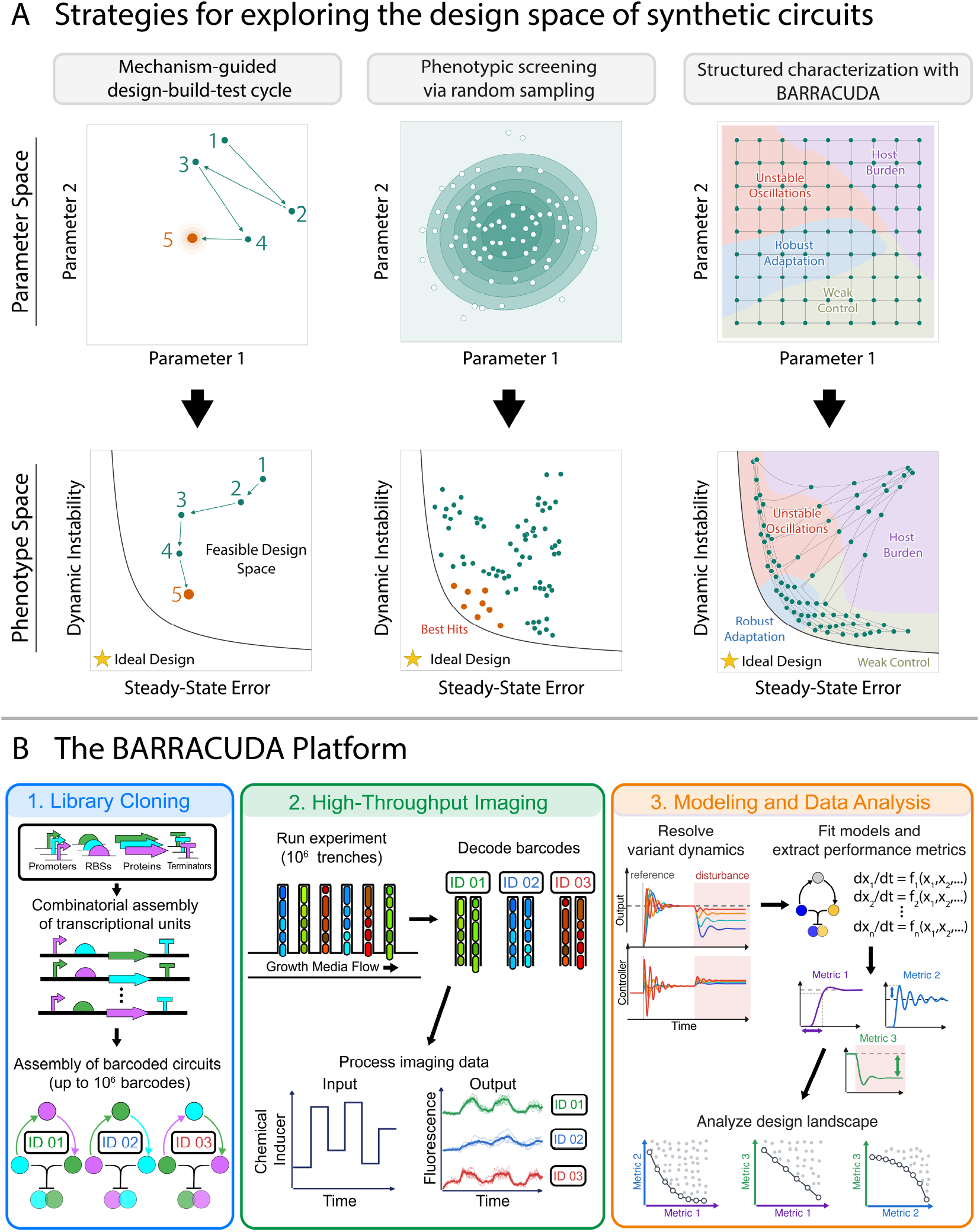
BARRACUDA enables structured, model-driven exploration of synthetic-circuit design spaces. **a**, Conceptual comparison of strategies for circuit engineering. The upper row shows designs sampled in parameter space and the lower row shows their corresponding locations in phenotype space, using steady-state error and dynamic instability as illustrative examples of performance measures. Left, mechanism-guided design–build–test cycles iteratively diagnose and improve individual designs but observe only a limited trajectory through parameter space. Center, phenotypic screening evaluates a broader sample and selects the best observed variants without necessarily inferring the structure of the intervening design landscape. Right, BARRACUDA systematically varies circuit parameters and uses genotype-resolved dynamic measurements to identify regions associated with distinct behaviors, including robust adaptation, weak control, unstable oscillations and host burden. The star denotes the ideal phenotype; orange points denote an improved design or selected hits. **b**, BARRACUDA workflow. Combinatorial assembly links each circuit to a genetic barcode. Long-read sequencing establishes a barcode–circuit lookup table, while mother-machine imaging records the dynamics of individual lineages under programmed chemical inputs. After imaging, sequential fluorescence in situ hybridization reads each lineage’s barcode; matching it to the lookup table assigns circuit identity to the recorded trajectory. These genotype-resolved measurements support mechanistic modeling, performance analysis and mapping of the design landscape.

Beyond identifying favorable variants, library-scale measurements can reveal predictive relationships between circuit composition and function. The CLASSIC platform ^18^ demonstrated this by combining flow cytometry and sequencing to profile endpoint expression across more than 10^5^ circuit designs in human cells. Neural networks trained on these measurements predicted unmeasured designs and revealed context-dependent rules for combining genetic parts, establishing an experimental basis for data-driven exploration of circuit design space.

Such approaches are powerful tools for mapping static input–output relationships, but regulatory circuits must also respond appropriately as conditions change. Endpoint measurements do not reveal whether regulation remains effective and stable during a perturbation. For example, an oscillating circuit and one exhibiting irregular fluctuations can produce indistinguishable fluorescence distributions despite their distinct temporal behavior. Dynamic responses also help distinguish mechanisms: different production, removal and interaction rates can produce similar endpoints but different transient responses to changing inputs.

To understand how circuit composition shapes these dynamic responses, we developed BARRACUDA (Barcoded ARRAys of Circuits Under Dynamic Actuation), a platform combining structured combinatorial libraries with multichannel time-lapse microscopy under programmed perturbations (Fig. 1a, right). Pooled libraries are grown in mother-machine microfluidic devices, where approximately one million spatially separated clonal lineages can be imaged in parallel. This supports experiments spanning 100 generations or more and sampling tens of thousands of circuit variants. Multiple fluorescent reporters track gene-expression dynamics and host responses in living cells. Each circuit design is represented by many lineages in the mother machine, providing extensive replication within the same experiment.

Building on the high-throughput imaging and optical pooled screening workflow developed for the MARLIN platform ^19^, BARRACUDA integrates combinatorial assembly with programmable environmental inputs and dynamical analysis (Fig. 1b). Each circuit carries a DNA barcode, and long-read sequencing reads barcode and circuit sequence together to establish their correspondence. After time-lapse imaging, barcode transcription is induced and the resulting RNA is decoded by sequential fluorescence in situ hybridization, linking each lineage’s recorded dynamics to its circuit design.

At each stage, we systematically vary components and expression levels to ask whether a shared mechanistic model can explain measurements across the entire circuit library. We then use the characterized components to build more complex circuits, asking whether their measured properties still explain circuit behavior when new interactions are introduced. Where predictions fail systematically, the discrepancies guide further experiments, model revision or component redesign.

We chose antithetic integral feedback (AIF) as a simple model to investigate how a regulatory principle depends on its molecular implementation. Maintaining an output at a desired level despite changing conditions is a longstanding goal of biological control ^5,20^. Integral feedback aims to precisely restore that output after a sustained disturbance, rather than merely reduce the disturbance’s effect ^21^. AIF provides a molecular architecture for this behavior: in an ideal stable regime, theory predicts adaptation despite sustained disturbances and changes in specified process parameters ^22^. Experimental implementations have demonstrated tunable regulation and disturbance rejection ^23,24^, but how much of the predicted robustness survives across components, expression levels and operating conditions remains a quantitative question. Establishing this robustness requires both applying external disturbances and systematically varying circuit parameters, distinguishing changes that feedback is predicted to compensate from those expected to shift the steady-state output.

To implement this circuit architecture we used a combination of sigma factors, which activate transcription, and antisigma factors, which bind and inhibit them. This pairing provides both transcriptional regulation and the protein sequestration needed to construct the feedback circuit. Using wild-type sigma–antisigma proteins expressed heterologously in *Escherichia coli*, we built libraries ranging from individual regulators and their inhibitory interactions to complete feedback circuits. We fitted mechanistic models to measurements from the simpler libraries to estimate component activities and interaction kinetics. We then used these estimates along with mathematical tools from control theory to derive predictions about the robustness of the assembled feedback circuits.

Unexpected feedback behavior motivated investigation of interactions between the introduced proteins and the host, and the design of de novo binders that retained sigma inhibition with reduced host-stress responses. Subsequent kinetic characterization revealed faster inferred removal of native-derived proteins than of most designed binders. Replacing the wild-type inhibitors with designed binders mitigated the anomalous overadaptation observed in the original circuits, while revealing a broader landscape of strong regulation and sustained oscillations. Counterintuitively, disturbances sometimes improved compensation rather than compromising it, suggesting nonlinear changes in how strongly feedback was engaged.

Together, these results establish a hierarchical approach to circuit engineering in which quantitative measurements transfer across design scales, discrepancies expose biological interactions missing from idealized models, and the resulting knowledge guides the construction of progressively more sophisticated gene-regulatory networks.

## 2 BARRACUDA enables quantitative characterization of combinatorial circuit libraries

Measuring many circuit designs in a pooled experiment requires a way to identify which circuit produced each recorded trajectory. To make this connection, BARRACUDA links each assembled circuit to a heritable DNA barcode that can be decoded via optical pooled screening after imaging, following the strategy established by the MARLIN protocol ^19^. Long-read sequencing establishes which circuit variant each barcode belongs to by capturing both sequences in the same read. Each circuit variant is associated with many distinct barcodes, providing independently sampled clones of that design.

The pooled library is loaded into a mother-machine microfluidic device, where individual clonal lineages occupy separate, spatially indexed growth trenches. We record their fluorescence dynamics under programmed inputs before reading their identities. After phenotypic imaging, barcode transcription is induced and the resulting RNA is decoded through sequential rounds of multiplexed error-robust fluorescence in situ hybridization (MERFISH). Matching this optical barcode to the sequencing-derived codebook identifies the circuit in each trench and assigns that identity to its recorded trajectory. Barcode readout is separate from the fluorescent reporters used to measure circuit activity. BARRACUDA combines this established genotype-resolved imaging workflow with rapid combinatorial library assembly, a sequencing-processing pipeline tailored to large combinatorial libraries, and automated input control ^25^, enabling structured characterization of circuit responses across design space.

In the cloning workflow, termed Checkpoint Cloning, unbarcoded circuits are assembled by multipart Golden Gate assembly, inspired by 3G Assembly ^26^. Full-length products are isolated by size before the 30-bit barcode cassettes are added in a final two-part assembly. Circuit libraries can be cloned in two days, with long-read sequencing yielding the codebook over the following 3–4 days. Imaging follows approximately one million trenches at 15-min resolution, and endpoint optical decoding typically identifies 400,000–800,000 trenches whose barcodes map to valid circuit designs.

Before constructing interacting regulatory networks, we began with the single transcriptional unit (TU), in which a promoter, ribosome-binding site (RBS), and coding sequence jointly determine gene expression. A TU is simple enough to admit a compact compositional model, yet it contains the same fundamental operations (input regulation, protein production, and measurement) that underlie more complex circuits. These calibrations are also needed to test feedback quantitatively: changing an RBS sequence does not by itself establish how much production has changed. Measuring the effective activities of the expression ladder supplies the parameter axis against which circuit responses can later be compared. We therefore used combinatorial TU libraries to test whether BARRACUDA measurements support predictive modeling across many combinations of the same parts and to establish a calibrated toolkit for subsequent circuit construction. Specifically, we sought inducible promoters whose responses could be parameterized as simple input–output relationships, a modular RBS ladder that varies production over a broad range with minimal context dependence, and bright fluorescent proteins that can be used to report on the resulting output dynamics.

To identify a suitable translation-rate ladder, we first screened 26 RBSs spanning three design families: the original set of bicistronic design (BCD) translational control elements ^8^, a recent redesign of the BCD RBSs ^27^, and a monocistronic RBS series ^28^ (Supplementary Fig. S1). The recently redesigned 12-member BCD series combined a broad, distributed expression range with the closest agreement to a separable expression model across reporter contexts (Supplementary Fig. S2), and we refer to it throughout as the BCD ladder. We retain the published RBS identifiers as labels; quantitative analyses use calibrated activities rather than the nominal design strengths. We therefore focused subsequent analysis on the subset containing 12 constitutive promoters ^29^, the 12-member BCD ladder and four fluorescent reporter proteins ^30,31^ (Fig. 2a). Of the 576 possible transcriptional units, 563 were recovered and measured. We modeled the fluorescence of promoter *i*, RBS *j* and reporter *k* as

**Figure 2.**
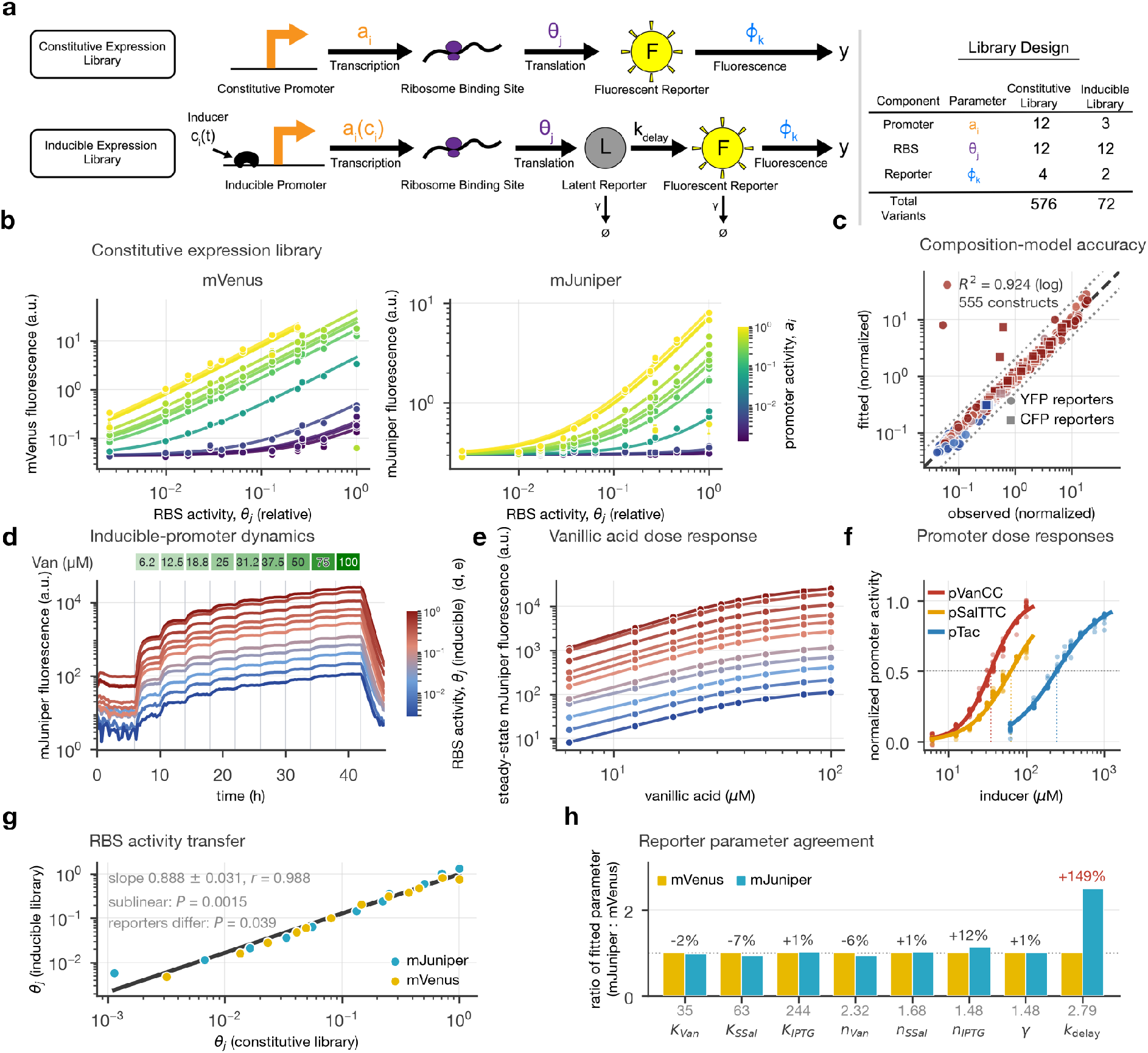
Combinatorial expression libraries calibrate modular components and inducible responses. **a**, Constitutive and inducible reporter-expression circuits and library designs, mapping promoter activity *a*_*i*_ or *a*_*i*_(*c*_*i*_), RBS activity *θ*_*j*_ and reporter gain *ϕ*_*k*_ onto transcription, translation and fluorescence measurement. The libraries contain 576 and 72 designs, respectively. **b**, Constitutive library comprising 12 promoters, the 12-member BCD ladder and four reporters; mVenus and mJuniper are shown. Points are construct measurements and curves are the joint model plotted against inferred RBS activity *θ*_*j*_ and colored by constitutive promoter activity *a*_*i*_. **c**, Model fit across 555 constructs (*R*^2^ = 0.92, log space; 576 designed and 563 measured). Dashed and dotted lines denote identity and twofold deviation. **d**, Median mJuniper trajectories for pVanCC circuits with BCD RBS variants during nine 4-h vanillic-acid plateaus (6.25–100 *µ*M) and washout. Color denotes *θ*_*j*_ inferred independently from the inducible library; its scale is therefore distinct from **b**. The BCD-0.1k–mJuniper combination showed no detectable induction and was excluded from the inducible calibration, leaving 11 mJuniper RBS variants. **e**, Steady-state values from **d** with affine variant-level fits sharing the half-maximal concentration *K* and Hill coefficient *n*. **f**, Dose responses of pVanCC, pSalTTC and pTac after two-parameter affine normalization of each variant. The collapse tests shared (*K, n*), not parameter-free prediction. **g**, Inducible versus constitutive *θ*_*j*_ for mVenus and mJuniper (*n* = 23). A common fit gives slope 0.888 *±* 0.031 and *r* = 0.988, sublinear with *P* = 0.0015. Because each reporter’s axis is normalized at unit activity, both intercepts are pinned to zero; on that parameterization a reporter-specific slope is supported (*P* = 0.039; 0.933 for mVenus, 0.859 for mJuniper). **h**, Ratios of parameters fitted independently through mJuniper and mVenus. Estimates of *K, n* and reporter loss *γ* agree within 13% (*γ*, 0.7%), whereas the reporter-dependent effective delay rate *k*_delay_ differs by 149%.

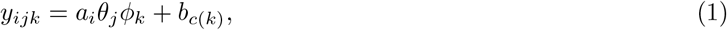

where *y*_*ijk*_ is measured fluorescence, *a*_*i*_ is promoter activity, *θ*_*j*_ is the calibrated relative activity of RBS *j, ϕ*_*k*_ is the fluorescence gain of reporter *k* and *b*_*c*(*k*)_ is the background in its fluorescence channel *c*(*k*). Fitting this model jointly across all four reporters and the complete promoter series gave *R*^2^ = 0.92 in log fluorescence space (Fig. 2b,c). Because each part appeared in many promoter–RBS–reporter combinations, this agreement tested whether a shared activity could describe its behavior across those contexts. The inferred promoter and RBS activities spanned 801-fold and 392-fold, respectively.

We next combined the 12-member BCD ladder with three small molecule-inducible promoters ^32^ (pVanCC, pSalTTC and pTac, regulated by VanR, NahR and LacI, respectively) and two bright fluorescent reporters (mVenus ^30^ and mJuniper ^31^) in a 72-member library (Fig. 2a). Automated actuation exposed the pooled library to successive inducer steps while preserving the complete response trajectory of each lineage (Fig. 2d). Rather than reducing these trajectories to plateau measurements, we fitted them directly with a two-state model that captures the delay between reporter production and observed fluorescence.

The model combines an inducible dose response with basal expression, protein removal and a delay between production and fluorescence. We fitted RBS activities across all three promoters within each reporter, while allowing promoter-specific dose responses and fitting mVenus and mJuniper independently (Supplementary computational methods, Eqs. S2 and S3). The delay is phenomenological rather than a direct measurement of fluorophore maturation. The resulting model captured the expression scale set by each RBS and the common dose-response shape of each promoter across the library (Fig. 2e,f). For visualization, plateau measurements were normalized by a variant-specific offset and scale while retaining the dynamically inferred dose-response parameters. Their collapse therefore tests a common response shape rather than constituting a parameter-free prediction.

The RBS activities inferred from the dynamic fits closely tracked those inferred independently from the constitutive library across both reporters (*r* = 0.988; Fig. 2g). The relationship was slightly sublinear (log–log slope 0.89), indicating modest compression of the expression ladder in the inducible context. The calibrated activities therefore transferred closely, but not exactly, between expression contexts.

Independent fits through mVenus and mJuniper also recovered similar promoter dose-response parameters and protein loss rates (Fig. 2h). The inferred loss rates corresponded to a half-life of approximately 28 minutes, closely matching the reported division time of our background strain in rich medium at 37 °C ^32^. By contrast, the effective delay between reporter production and observed fluorescence differed substantially between reporters. Together, these libraries show that BARRACUDA can support quantitative modeling at scale while testing component properties across constructs, reporters, and experimental contexts. Combinatorial fitting tests whether shared part activities can explain many different constructs. Independent fits through different reporters test whether the same regulatory parameters can be recovered using different measurement proteins, while comparison of constitutive and inducible libraries tests how well expression activities transfer between contexts. The observed agreement, together with the context-dependent compression and reporter-specific delays, provides a quantitative basis for deciding which properties can be reused and which require separate calibration. These first libraries therefore establish both a characterized expression toolkit and a strategy for testing the mechanistic assumptions carried into more complex circuits.

## 3 Combinatorial screening identifies compatible regulators for feed-back control

Having established a calibrated expression ladder and quantitative control through inducible promoters, we next sought compatible regulatory proteins from which to build an AIF circuit ^22^. Our design calls for modular regulators that can drive the production of downstream components. We chose to implement these functions through transcriptional activation, using two regulators that could operate simultaneously. One provides the activity we aim to regulate (the process) while the other drives its production as part of the controller. A feedback inhibitor must respond to process activity and sequester the controller, while a separately induced inhibitor provides a disturbance to the process. Each regulator must therefore activate a distinct transcriptional output, be efficiently sequestered by its cognate inhibitor and remain tunable across a broad expression range. Extracytoplasmic-function (ECF) sigma factors and their antisigma factors provide a compact molecular implementation of these requirements ^33,34^. Sigma factors associate with RNA polymerase and direct transcription from cognate promoters, whereas antisigma factors bind and inactivate their partner through molecular sequestration. In principle, this combines positive transcriptional regulation with a specific negative interaction suitable for antithetic feedback. In practice, however, performance at a single expression level does not establish that a sigma–antisigma pair will be useful as a control module. Two systems must remain mutually orthogonal, retain broad and independently tunable activity ranges, provide sufficient capacity to inhibit their cognate sigma factors, and remain functional at the expression levels required by the closed-loop circuit.

We therefore screened five candidate systems across their expression and interaction contexts. We selected ECF11, ECF20, ECF22 and ECF27 from a previously characterized collection ^35^ of ECF sigma factors and included SigW, which had previously been used to implement antithetic integral feedback ^23^. We constructed an 18,000-member combinatorial library in which pVanCC expressed one of the five sigma factors through the 12-member BCD ladder, pSalTTC expressed one of the five antisigma factors through the same ladder, and one of five sigma-dependent promoters drove mJuniper (Fig. 3a). Successive vanillic acid (Van) and sodium salicylate (SSal) steps separately probed sigma activation and antisigma-mediated suppression within a single experiment. The combinatorial design paired every sigma factor with every reporter promoter and antisigma factor while independently varying sigma and antisigma expression. This allowed us to quantify activation and sequestration strength and distinguish cognate from non-cognate interactions.

**Figure 3.**
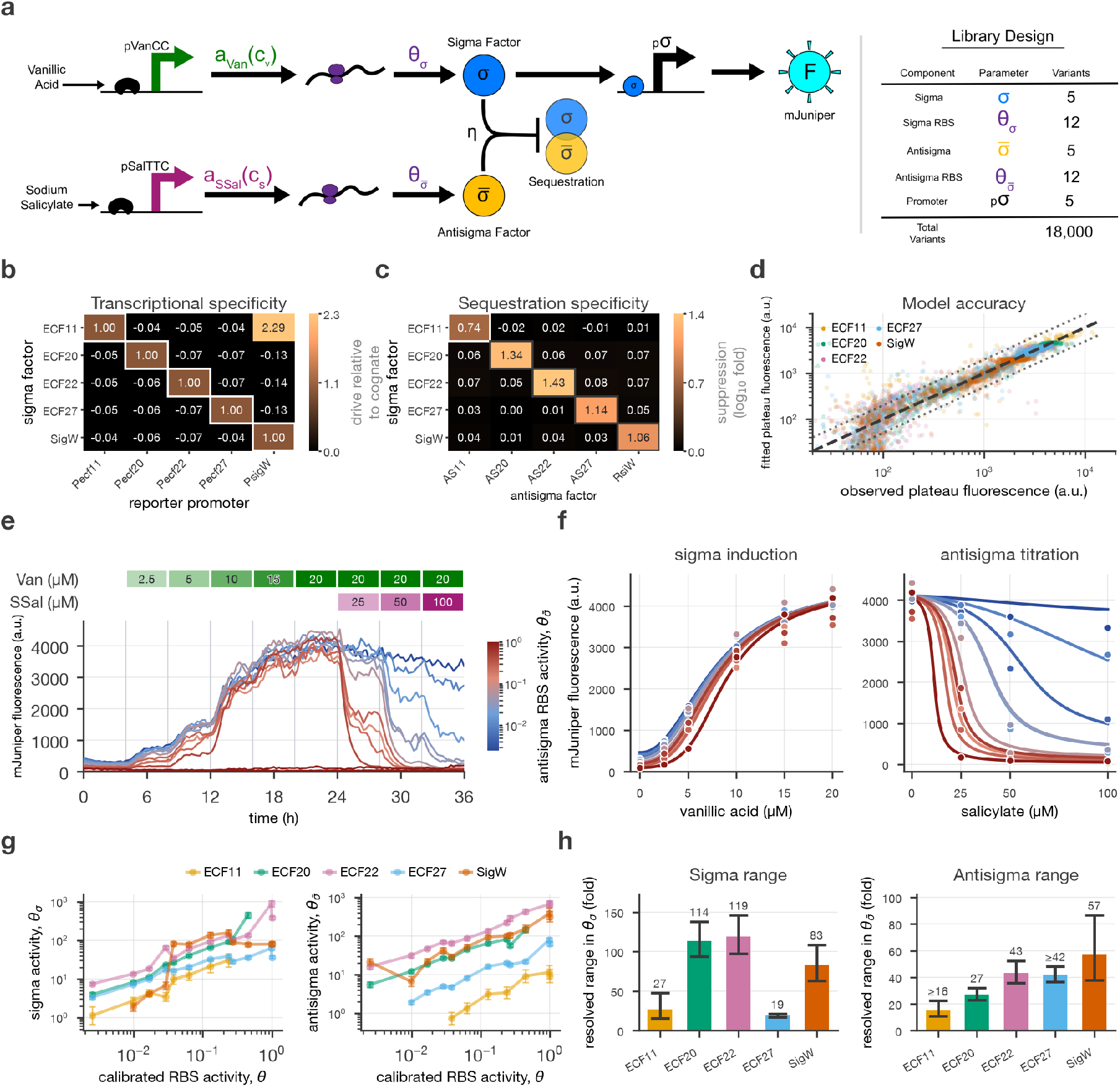
Dynamic screening identifies orthogonal and tunable sigma–antisigma systems. **a**, Design of the 18,000-member library comprising five sigma factors, 12 sigma RBSs, five antisigma factors, 12 antisigma RBSs and five reporter promoters. Vanillic acid induces sigma expression before salicylate titrates antisigma expression. **b**, Transcriptional specificity. Each column is normalized to the drive that promoter receives from its own cognate sigma factor, so the diagonal has a slope of 1 by construction. ECF11 activates *P*_SigW_ 2.29-fold more strongly than SigW does; the other 19 non-cognate interactions show no detectable activation. **c**, Sequestration specificity, reported as the median log_10_ suppression between 0 and 100 *µ*M salicylate at fixed sigma induction. Cognate suppression spans 0.74–1.43 decades. **d**, Cognate model fit for 4,272 steady-state plateaus. Each regulatory pair was fitted separately and all points are training data (*R*^2^ = 0.9312, linear space; 84.0% within twofold). **e**, Median ECF20–AS20 trajectories with the sigma RBS fixed and the 12-member AS20 RBS ladder varied. **f**, Observed plateaus and fitted responses for sigma induction (left) and antisigma titration (right) for the slice shown in **e. g**, Effective sigma and antisigma activities, *θ*_*σ*_ and 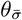, versus the reference RBS activity *θ*_*j*_. Error bars show 95% asymptotic Fisher intervals. **h**, Resolved fold ranges of the effective sigma and antisigma ladders. Error bars are propagated 95% Fisher intervals; inequalities denote conservative bounds after excluding insufficiently supported rungs.

To infer effective sigma and antisigma activities, we fitted the steady states of a model combining inducible production, dilution-dominated loss, irreversible sequestration and a saturating sigma-dependent reporter (Supplementary Eq. S5). Inducer-response parameters were inherited from the reporter screen, whereas expression and basal activities were estimated in each regulatory context. The fitted RBS activities, *θ*_*σ*_ and 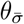, quantify production of the sigma and antisigma proteins. Because fluorescence does not establish absolute molecular concentrations, we compare their ordering and fold ranges within each system rather than absolute activities across different sigma-dependent reporters.

The screen first resolved regulatory specificity. Each promoter’s responses were normalized to the drive it received from its own cognate sigma factor. Nineteen of the twenty non-cognate sigma–promoter combinations produced no detectable activation, whereas ECF11 activated *P*_SigW_ 2.29-fold more strongly than SigW itself did (Fig. 3b). Relative to its own cognate promoter, ECF11 drove *P*_SigW_ 0.86-fold as strongly, so this cross-activation is substantial on the scale of the target promoter’s native input rather than of ECF11’s own output. Antisigma-mediated suppression was similarly specific across all five systems, reaching approximately 5.5–27-fold for cognate pairs while non-cognate effects remained near zero (Fig. 3c). Fits performed separately for each regulatory pair captured the cognate response surfaces across both expression axes. Across 4,272 fitted cognate plateaus, the model achieved *R*^2^ = 0.9312 in linear fluorescence space and placed 84.0% of observations within twofold of their fitted values (Fig. 3d). For example, the model reproduced both ECF20 induction and its graded suppression across the 12-member AS20 expression ladder (Fig. 3e,f).

ECF20 and ECF22 combined mutual specificity with broad, tunable expression ranges for both sigma and antisigma proteins, making them the most suitable pair for feedback construction (Fig. 3g,h). Effective activities generally increased with the reference RBS activity, but their usable ranges depended strongly on the regulatory protein. ECF27 compressed the sigma-expression range, while SigW produced similar activities across several nominally strong RBS variants, limiting the expression variation available from those parts. ECF11 showed substantial off-target promoter activation and poor library representation, with several nominally strong sigma-expression variants absent from the recovered data. Given these results, we selected ECF20 and ECF22, together with their cognate antisigma factors, as the core regulatory components of an AIF network. We then asked whether the specificity and tunability observed in open loop would translate into predictable closed-loop dynamics.

## 4 Feedback circuits compensate for perturbations but reveal unexpected dynamics

Building on this open-loop characterization, we next assembled the selected sigma–antisigma systems into a closed-loop AIF network. We evaluated its performance by measuring both process and controller activity while varying expression levels and applying external disturbances. Tools from control theory provide a quantitative framework for interpreting these measurements ^36–40^. In an ideal stable AIF regime, the process should maintain its set point despite changes in its production capacity or a sustained disturbance, while the controller changes to compensate ^22,23^. Measuring both activities therefore allows us to distinguish feedback-mediated robustness from a superficially flat output caused by reporter saturation or limited dynamic range.

For the assembled circuit, we denote the free controller sigma and its inhibitor by *c*_1_ and *c*_2_, and the process sigma and its disturbance inhibitor by *p* and *d*. Expression parameters retain the symbol *θ*, but their subscripts now identify circuit roles rather than individual RBS rungs (Supplementary Table S1).

We constructed an ECF20–ECF22 circuit in which vanillic acid drives chromosomally integrated ECF20, the controller sigma *c*_1_ (Fig. 4a). ECF20 activates expression of the process sigma ECF22, *p*, through an RBS with activity *θ*_*p*_. ECF22 in turn drives AS20, *c*_2_, the cognate antisigma factor of ECF20, through an RBS with activity *θ*_*c*_, closing the negative-feedback loop through sequestration of *c*_1_. Salicylate independently induces the anti-ECF22 disturbance AS22, *d*, through an RBS with activity *θ*_*d*_. Chromosomal *P*_ECF20_–mJuniper and *P*_ECF22_–mVenus reporters measure controller and process activity, respectively. Each of the three plasmid-borne transcriptional units samples the 12-member BCD ladder, producing 12^3^ = 1,728 parameter combinations, of which 1,723 (99.7%) were recovered in the closed-loop configuration and sampled by 320,451 decoded lineages (median 86 lineages per combination); the sensitivity analyses below use the 1,655 combinations retaining at least ten lineages.

**Figure 4.**
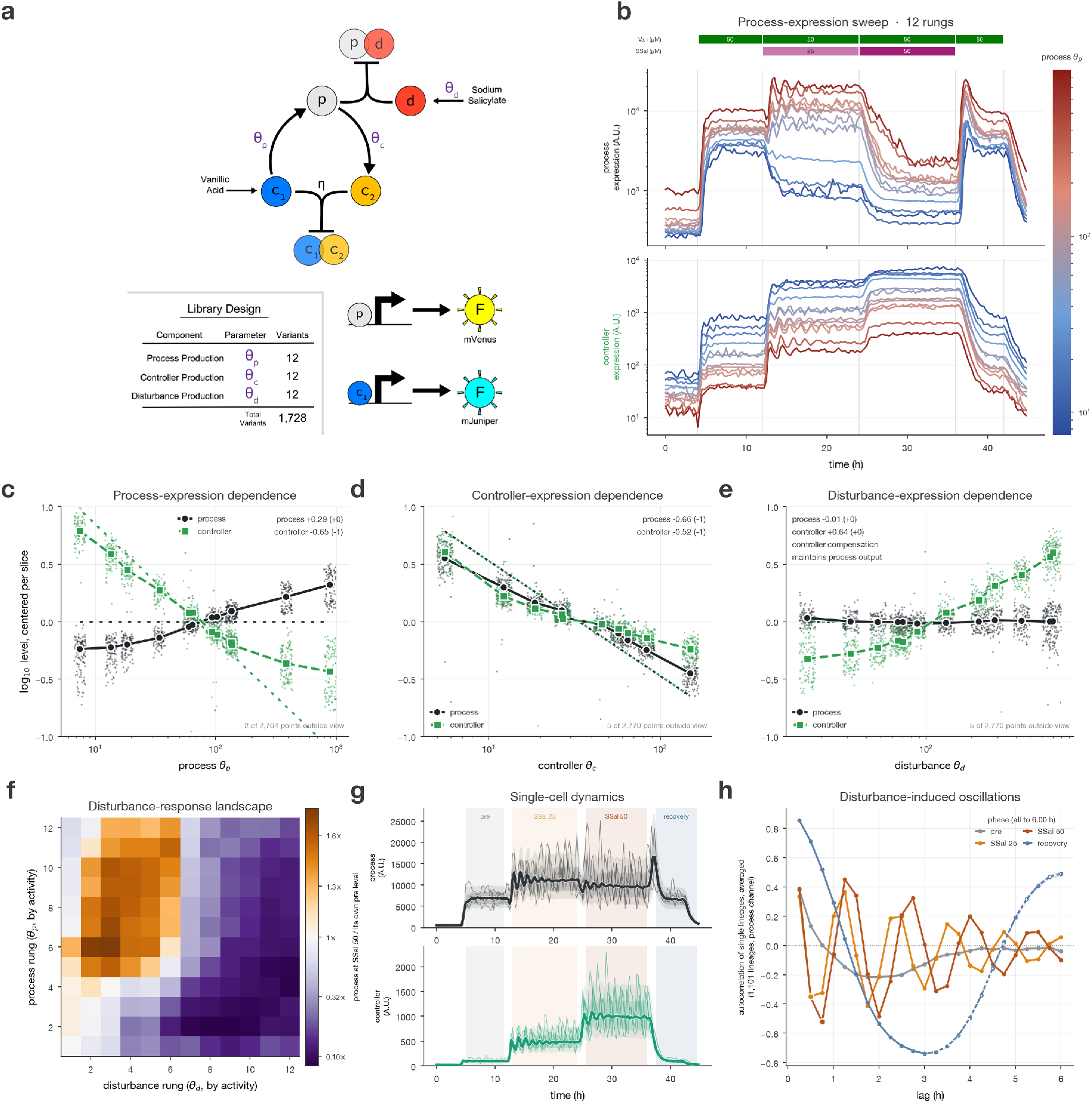
An antithetic integral control circuit library reveals disturbance rejection, overadaptation and oscillations. **a**, Circuit and library architecture. Chromosomal ECF20 (*c*_1_) drives plasmid-encoded ECF22 (*p*) through *θ*_*p*_; ECF22 drives AS20 (*c*_2_) through *θ*_*c*_; and salicylate induces AS22 (*d*) through *θ*_*d*_. The sequestering pairs are (*c*_1_, *c*_2_) and (*p, d*). Each RBS samples the 12-member BCD ladder. **b**, Median process-reporter (mVenus) and controller-reporter (mJuniper) trajectories while sweeping *θ*_*p*_, with *θ*_*c*_ and *θ*_*d*_ fixed. Lines aggregate all decoded trenches assigned to each genetic variant. **c–e**, Slice-centered log_10_ process and controller levels as functions of *θ*_*p*_, *θ*_*c*_ and *θ*_*d*_, respectively, before disturbance induction. Small points show individual fixed-background slices; large symbols and connecting lines summarize each rung. Reported elasticities are medians of independent within-slice ordinary least-squares slopes (119, 143 and 118 slices), not slopes fitted to the pooled points. Low-signal genotypes are excluded; no predisturbance genotype is removed by this criterion in these data. Black solid/green dashed lines denote process/controller. Dotted lines and parenthetical values show the zero-basal-disturbance ideal-AIF reference; nonzero basal disturbance changes the controller expectations (Supplementary Eq. S26). **f**, Disturbance-response landscape across all 12 process-expression and 12 disturbance-expression rungs. Color indicates the median per-variant process-output ratio at 50 *µ*M salicylate relative to predisturbance output, pooling the 12 controller-expression rungs; cells require at least three variants. Controller-expression dependence is resolved separately in Section 7. **g**, Process and controller dynamics for a well-sampled variant exhibiting sustained overadaptation and disturbance-induced single-lineage oscillations. Thick lines denote medians, nested shaded bands denote interquartile and 10th–90th percentile ranges, and thin lines show three representative lineages. **h**, Mean per-lineage autocorrelation of detrended process trajectories before disturbance, during 25 and 50 *µ*M salicylate, and during recovery; shading denotes one standard error. The broad recovery trough and rebound are not interpreted as an oscillation period.

The feedback acts through the production of the controller inhibitor. If process activity falls, production of *c*_2_ falls, releasing more free *c*_1_ to drive process expression. If process activity rises, increased *c*_2_ production restrains *c*_1_. Here, the regulated output is ECF22 activity, measured through its transcriptional reporter.

To define the ideal behaviors against which this library could be compared, we described the circuit with a four-state irreversible-sequestration model,

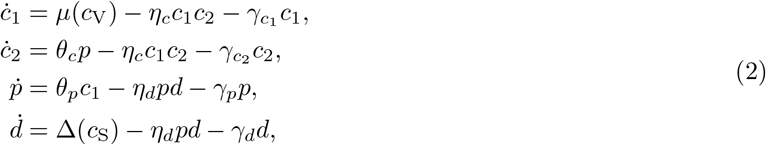

where *µ* is the vanillic acid-dependent controller input and Δ is the salicylate-dependent disturbance input, including basal expression in the absence of salicylate. The parameters *η*_*c*_ and *η*_*d*_ describe sequestration on the controller and disturbance branches; the *γ* terms describe independent removal by dilution or degradation. The model and its limits are developed in the Supplementary feedback theory.

In the ideal AIF limit, both controller species are removed only through mutual sequestration (i.e., 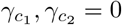. Their shared removal term cancels when the controller equations are subtracted, so the difference *c*_1_ − *c*_2_ accumulates the mismatch between the imposed input *µ* and the feedback production *θ*_*c*_*p*. This molecular balance stores accumulated output error. At a stable steady state, the mismatch vanishes, giving

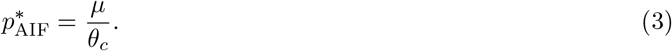

The process set point is therefore independent of process expression *θ*_*p*_ and disturbance expression *θ*_*d*_, while scaling inversely with controller expression *θ*_*c*_. Independent dilution or degradation of the controller species changes this balance, making integration leaky and limiting the disturbance load the circuit can oppose ^37,38^. The controller steady state and explicit fast-sequestration predictions are developed in Supplementary Eqs. S20–S24.

We quantified these responses using elasticities, which measure the fractional change in an output produced by a fractional change in an expression parameter ^41–44^:

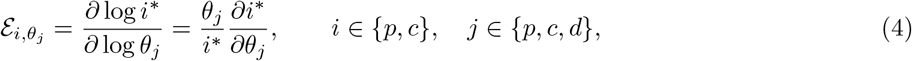

Here, the first index identifies process or controller output, with *c* referring to *c*_1_, and the second identifies the varied expression parameter. *ℰ* = 0 denotes perfect parameter insensitivity, and *ℰ* = 1 denotes a proportional response. Calibrated RBS activities supply the parameter axis, allowing the theoretical predictions to be tested by varying one expression ladder at a time.

Ideal feedback predicts zero process elasticity to both process and disturbance expression, with compensatory changes in the controller. The expected controller responses depend on the disturbance load, which can be nonzero even before salicylate addition because of basal production of the disturbance inhibitor. This basal load changes the predicted controller elasticities without changing the ideal process insensitivity. The complete ideal and leaky predictions are given in Supplementary Eqs. S25 and S26.

Experimental elasticities were estimated from fluorescence after subtracting a locally estimated additive background, calibrated against cells lacking the corresponding fluorescent-protein gene. Constant multiplicative measurement scales cancel from an elasticity, but interpretation as a concentration response additionally requires an approximately proportional reporter response over the analyzed range.

To test these predictions, we held two RBSs fixed while sweeping the third across the BCD ladder and measured phase-level reporter output before disturbance induction (50 *µ*M vanillic acid, 0 *µ*M salicylate). We estimated a log–log slope independently within each fixed-background slice, excluding low-signal rungs before fitting, and then summarized the qualifying slopes by their median (Supplementary computational methods). Increasing process expression *θ*_*p*_ produced only a modest change in process output but a strong compensatory decrease in controller activity (Fig. 4b,c). The corresponding summary elasticities, 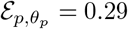 and 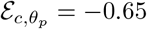, lie between the open-loop limit, in which the process absorbs the full change in its own expression and the controller does not respond (1 and 0), and the ideal-feedback limit, in which that response is transferred entirely to the controller (0 and 1). The curvature across the ladder further indicates that the effective response changes across the sampled operating range, rather than being described by one local elasticity.

Increasing controller expression *θ*_*c*_ decreased both process and controller outputs, with summary elasticities 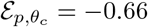 and 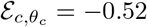 (Fig. 4d). The process response again lies between the weak-feedback limit of zero and the ideal value of 1. The controller response is compatible with the ideal prediction in the presence of basal disturbance, although variation among slices indicates that one common gain and disturbance load cannot describe the complete library (Supplementary Eq. S26).

The clearest evidence for active disturbance rejection came from the *θ*_*d*_ sweep. Process output remained nearly invariant across the disturbance-expression ladder 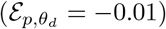, whereas controller activity increased strongly (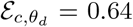 Fig. 4e). We note that these measurements still take place before salicylate is added to induce the disturbance, meaning that the controller is responding to leaky expression of AS22 from the uninduced pSalTTC promoter. Thus, the circuit maintains its process output by increasing controller action rather than through passive saturation of the reporter. The positive controller elasticity further implies nonzero basal disturbance load, as predicted by Supplementary Eq. S26. Together, the three parameter sweeps show the complementary process and controller responses expected from feedback while revealing systematic departures from the simplest ideal-AIF limit.

We next examined disturbance rejection under 50 *µ*M vanillic acid and 50 *µ*M salicylate induction across the full *θ*_*p*_-by-*θ*_*d*_ design space (Fig. 4f). At low-to-intermediate disturbance expression, broad regions of the library maintained or increased process output after AS22 induction. At higher *θ*_*d*_, however, performance collapsed sharply, consistent with the finite capacity to reject disturbances described in Supplementary Eq. S23. The location of this boundary shifted with *θ*_*p*_: stronger process expression increased the disturbance load that could be opposed before free ECF22 was depleted. The resulting landscape therefore contains both a broad disturbance-rejecting regime and a distinct capacity boundary rather than a single uniformly adaptive phenotype. Despite this evidence for feedback, the library also exhibited behaviors not accounted for by the fast-sequestration description ^37^. Many variants overadapted, reaching process levels above their predisturbance state and remaining elevated during sustained disturbance (Fig. 4g). In the leaky model, increasing the disturbance Δ can only reduce *p*^*^; in the ideal limit, it leaves *p*^*^ unchanged. Neither regime permits steady-state overadaptation. The same variants frequently developed pronounced single-lineage oscillations after disturbance induction. These oscillations were partially obscured by population-level aggregation but were reproducible across independently tracked lineages and disappeared during washout (Fig. 4g,h).

In the limit of fast sequestration kinetics, the reduced circuit dynamics are approximately linear ^37^. Within this regime, a constant disturbance shifts the equilibrium without changing the Jacobian or its eigenvalues, and therefore without changing stability. A circuit that is stable before disturbance consequently cannot develop sustained oscillations solely because that disturbance is applied. The model also imposes a separate steady-state constraint: even outside the fast-sequestration limit, increasing disturbance at fixed parameters can lower the equilibrium process output or leave it unchanged, but cannot raise it above its predisturbance value. Disturbance-dependent instability therefore lies beyond the linear reduction, whereas equilibrium overadaptation lies beyond the fixed-parameter four-state model (Supplementary feedback theory).

These discrepancies led us to consider interactions between the heterologous antisigma factors and the *E. coli* host. Native ECF antisigma factors contain transmembrane and hydrophobic regions that could engage membrane quality control or cytosolic proteostasis pathways. Such responses could increase degradation of free antisigma proteins, removing inhibitors without the mutual sequestration on which the feedback loop depends. If this degradation changed with antisigma expression, the circuit would acquire expression-dependent removal rates and additional host-mediated dynamics absent from the fixed-parameter model.

## 5 De novo binders preserve sigma inhibition with reduced stress responses

We designed a single multiplexed library to test whether native antisigma expression elicited host-stress responses and to screen alternative inhibitors for rebuilding the feedback circuit (Fig. 5a). The library combined native and truncated antisigma factors with de novo binders. Stress-responsive reporters assessed host effects, while measurements of sigma activity quantified inhibition and specificity, allowing us to investigate the proposed host interaction and evaluate potential replacements in the same experiment.

**Figure 5.**
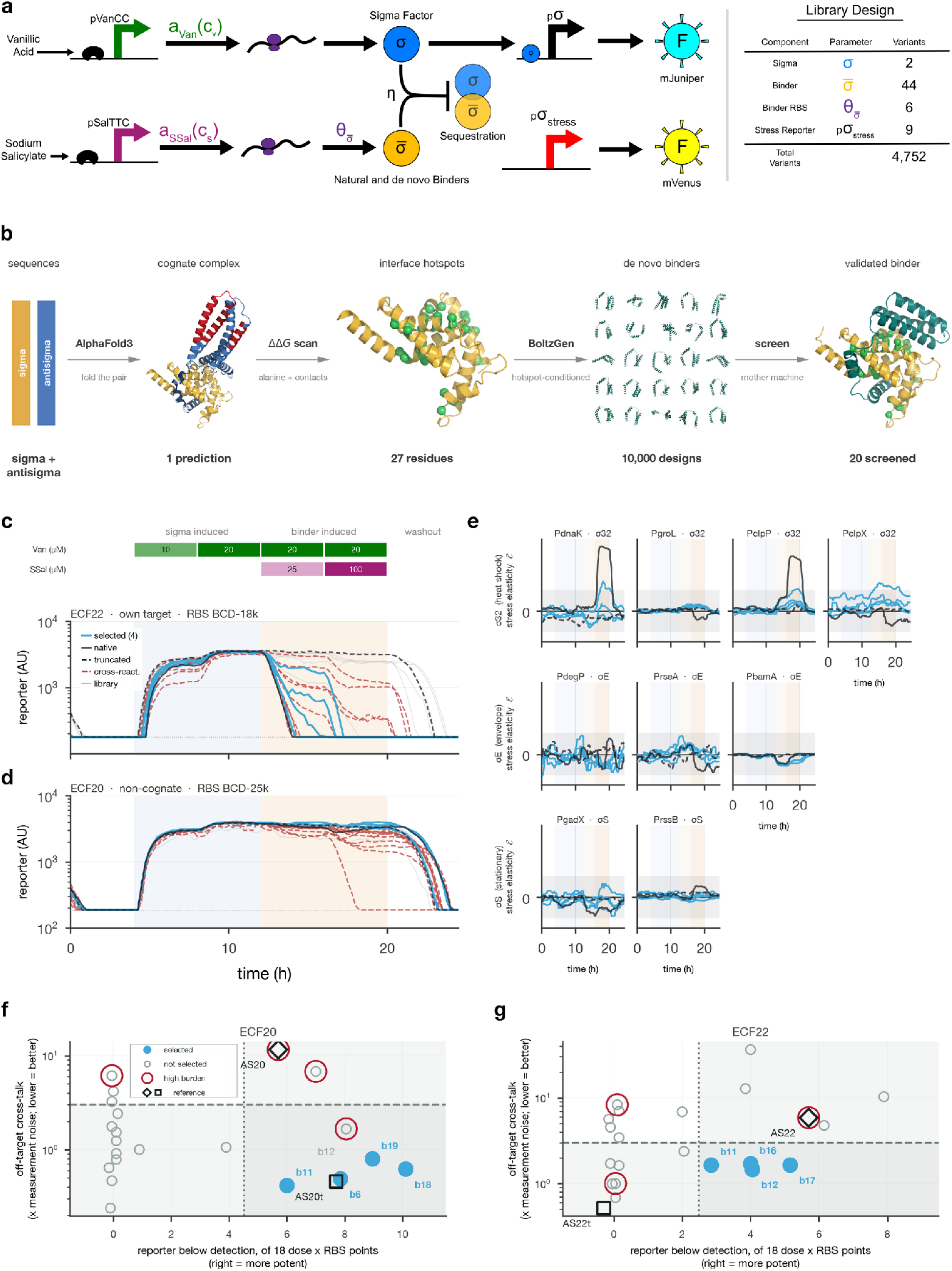
Selected de novo binders combine specific sigma inhibition with reduced stress responses. **a**, Single multiplexed library testing 44 binders against both sigma factors across six binder RBSs and nine stress reporters (4,752 designed variants). **b**, Design workflow. The two BoltzGen campaigns together generated approximately 10,000 designs per target; 20 binders per target were synthesized. Structures are AlphaFold3 predictions. **c,d**, Median cognate ECF22 (**c**, BCD-18k) and noncognate ECF20 (**d**, BCD-25k) reporter trajectories for ECF22 binders. Blue, selected; black solid/dashed, native/truncated AS22; gray, other designs. Dashed red denotes graded off-target suppression, distinct from the composite gate in **f,g. e**, Stress elasticities using all six RBSs and the shared calibration from Fig. 2 for every binder. Gray bands mark *±*0.1175, three pooled pre-induction standard deviations (5–11 h; Supplementary Fig. S4). Promoters are grouped by associated stress regulon. **f,g**, Cognate suppression versus cross-talk for all 20 ECF20 (**f**) and ECF22 (**g**) designs and native/truncated controls. Suppression counts the 18 RBS–salicylate combinations reaching the background-subtraction floor. Cross-talk detects graded suppression or rung-dependent changes in either direction (Supplementary computational methods). Blue, selected; red rings, burden exclusions; diamonds/squares, native/truncated controls. Dashed cutoffs (cross-talk below 3; at least 5 or 3 suppressed combinations, respectively) describe the selected sets retrospectively.

The replacement designs were motivated by the substantial regions of AS20 and AS22 outside their predicted sigma-binding interfaces. We designed compact binders targeting the corresponding surfaces on ECF20 and ECF22, aiming to preserve inhibition without reproducing the extended architecture of the native proteins. We used BoltzGen, a generative protein-design pipeline that proposes candidate binder sequences and structures for specified targets ^45^.

We performed two BoltzGen campaigns for each sigma-factor target ^45^. A sequence-conditioned campaign targeted residues identified by proximity and contact analysis of predicted native complexes. A structure-conditioned campaign instead supplied precomputed AlphaFold3 complexes ^46^ and targeted interface hotspots identified by Rosetta-based computational alanine scanning ^47,48^ and PLIP contact analysis ^49^. Together, the two campaigns generated approximately 10,000 designs per target. Designs were ranked using a composite BoltzGen interface score, after which the top-ranked subset for each target was re-evaluated with AlphaFold3. From these candidates, we synthesized 20 de novo binders per target, selected for both computational rank and structural diversity, together with the corresponding native and truncated antisigma controls (Fig. 5b). Binder identifiers were assigned independently for each target, so we write the target prefix explicitly; for example, ECF20-b11 and ECF22-b11 are unrelated proteins.

We analyzed all 44 binder genotypes against both sigma-factor systems in a single multiplexed library. In each system, vanillic acid induced the sigma factor, its cognate promoter drove mJuniper, and salicylate induced the test binder through one of six RBS parts sampled from the calibrated BCD ladder ^27^. Each binder was therefore tested for suppression of both its intended target and the other sigma factor. These dimensions defined 44 × 6 × 2 = 528 functional combinations, each additionally crossed with one of nine stress-responsive promoters, giving 4,752 designed variants, of which 4,652 were recovered and measured (Fig. 5a).

Several ECF22 binders suppressed the cognate reporter as strongly as native AS22, but some also altered the noncognate ECF20 reporter (Fig. 5c,d; corresponding ECF20 measurements in Supplementary Fig. S3). We quantified these off-target responses relative to pre-induction variation, detecting both suppression and expression-dependent changes in the opposite direction. All slope-based comparisons used the shared six-rung calibration from Fig. 2. An operational specificity criterion retained 16 of 20 ECF20 binders and 10 of 20 ECF22 binders (Fig. 5f,g; Supplementary computational methods).

We also screened for nonspecific burden using expression-dependent changes in a constitutive chromosomal mKate2Hyb reporter. This proxy excluded one otherwise specificity-passing binder per target. Native AS20 and AS22 exceeded both screening cutoffs, but their off-target responses differed: AS22 suppressed the noncognate reporter, whereas AS20 increased it and also altered the constitutive reporter. AS20’s response was therefore consistent with host-mediated effects rather than off-target suppression; the reporter statistics alone do not establish direct cross-binding.

Among designs with low off-target effects and burden, we compared how broadly each binder suppressed its cognate reporter across the sampled expression and inducer conditions (Fig. 5f,g). We carried forward four binders for ECF20 (ECF20-b6, ECF20-b11, ECF20-b18 and ECF20-b19) and four for ECF22 (ECF22-b11, ECF22-b12, ECF22-b16 and ECF22-b17). Selection combined specificity, burden and cognate suppression, independently of the stress-reporter measurements and subsequent kinetic fits.

We next used the stress-responsive reporter carried by each variant to ask whether native antisigma factors affected host physiology and whether the designed binders altered these effects. The nine promoters sampled several native *E. coli* stress programs, including the RpoH cytoplasmic protein-folding response, the RpoE envelope-stress response and the RpoS general stress response ^33,50,51^. At each time point, the six RBS variants allowed us to estimate a stress elasticity,

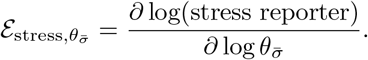

Here, 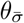 denotes the reference activity *θ*_*j*_ of the RBS driving binder expression, calibrated in the constitutive-expression library (Fig. 2), rather than a measured protein abundance or a binder-specific kinetic parameter. The same six-rung axis is used for every binder and both targets, without requiring the later kinetic fits. These slopes distinguish responses that depend on binder expression from changes shared across the RBS series, assuming that the calibrated relative activities transfer to the binder-expression context. A flat elasticity therefore does not exclude a stress response shared across all expression levels.

Native AS22 produced pronounced expression-dependent responses from *P*_dnaK_ and *P*_clpP_, both members of the RpoH heat-shock regulon ^50^ (Fig. 5e). The response was promoter-specific rather than regulon-wide: *P*_clpX_ and *P*_groL_ did not show comparably sustained positive responses. This pattern was consistent with cytoplasmic protein-folding stress, although promoter activity alone does not establish protein localization or degradation route ^52^. Native AS20 instead most strongly activated *P*_degP_, a member of the RpoE envelope-stress regulon ^33^ (Supplementary Fig. S3c). The other envelope-stress reporters did not show the same response: *P*_bamA_ had little expression-dependent signal, and *P*_rseA_ responded at about the level of the designed binders. The largest post-induction stress elasticities were 0.59 for AS20 and 0.36 for AS22, exceeding those of the selected binders against the corresponding target. AS20 truncation strongly reduced the response, leaving a brief, small excursion above the reference threshold.

To compare response magnitude and persistence, we integrated positive elasticity above the shared reference threshold from salicylate addition onward. The integrals were 2.52 and 0.94 h for AS20 and AS22, approximately 60.8- and 6.5-fold greater than the largest integrals among the selected binders against each target, ECF20-b6 and ECF22-b11, respectively (Supplementary Fig. S4). Two of the four selected binders per target exceeded the threshold, but their integrated responses were substantially smaller than those of the native antisigma factors. The designed binders therefore greatly reduced these expression-dependent host-stress reporter responses.

Native antisigma factors elicited substantially stronger host-stress responses than the selected de novo binders. However, these responses could occur without changing the proteins’ kinetics, leaving open whether the native proteins exhibited the enhanced turnover proposed to compromise feedback.

## 6 Optimal experimental design resolves open-loop component kinetic parameters

Before rebuilding the feedback circuit with the selected binders, we needed to characterize the kinetics of both native and designed components. This would supply production, sequestration and removal parameters for circuit design, while testing whether the native antisigma factors exhibited the enhanced turnover proposed to contribute to the original feedback anomalies.

Turnover was particularly important because dilution or degradation can remove an antisigma factor without matching removal of its sigma partner, making integration leaky. The stress responses raised a further possibility: antisigma expression might alter host protease activity, so that inhibitor degradation varies with induction level. By altering the antisigma available to sequester sigma, this coupling could shift steady-state regulation and introduce additional host-mediated dynamics, potentially contributing to the overadaptation and oscillations observed in the first feedback library.

To test the turnover hypothesis, we needed to distinguish each binder’s independent removal rate, 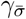, from its effective sequestration rate, *η*, while calibrating production across the expression ladders. Because different combinations of these properties could produce similar suppression, we sought dynamic perturbations that would make the alternative kinetic explanations produce distinguishable trajectories.

We constructed open-loop characterization libraries for 11 sigma–antisigma architectures: six based on ECF20 and five based on ECF22 (Fig. 6). Each architecture combined a 12-member *σ* RBS ladder with a 12-member antisigma RBS ladder, sampling the native antisigma factors, truncated AS20 (AS20t) and the de novo binders selected in Fig. 5. We described each architecture using a four-state dynamic model comprising *σ*, antisigma and two reporter states, the second of which is observed as fluorescence with an additive background, with irreversible bimolecular sequestration between *σ* and antisigma.

**Figure 6.**
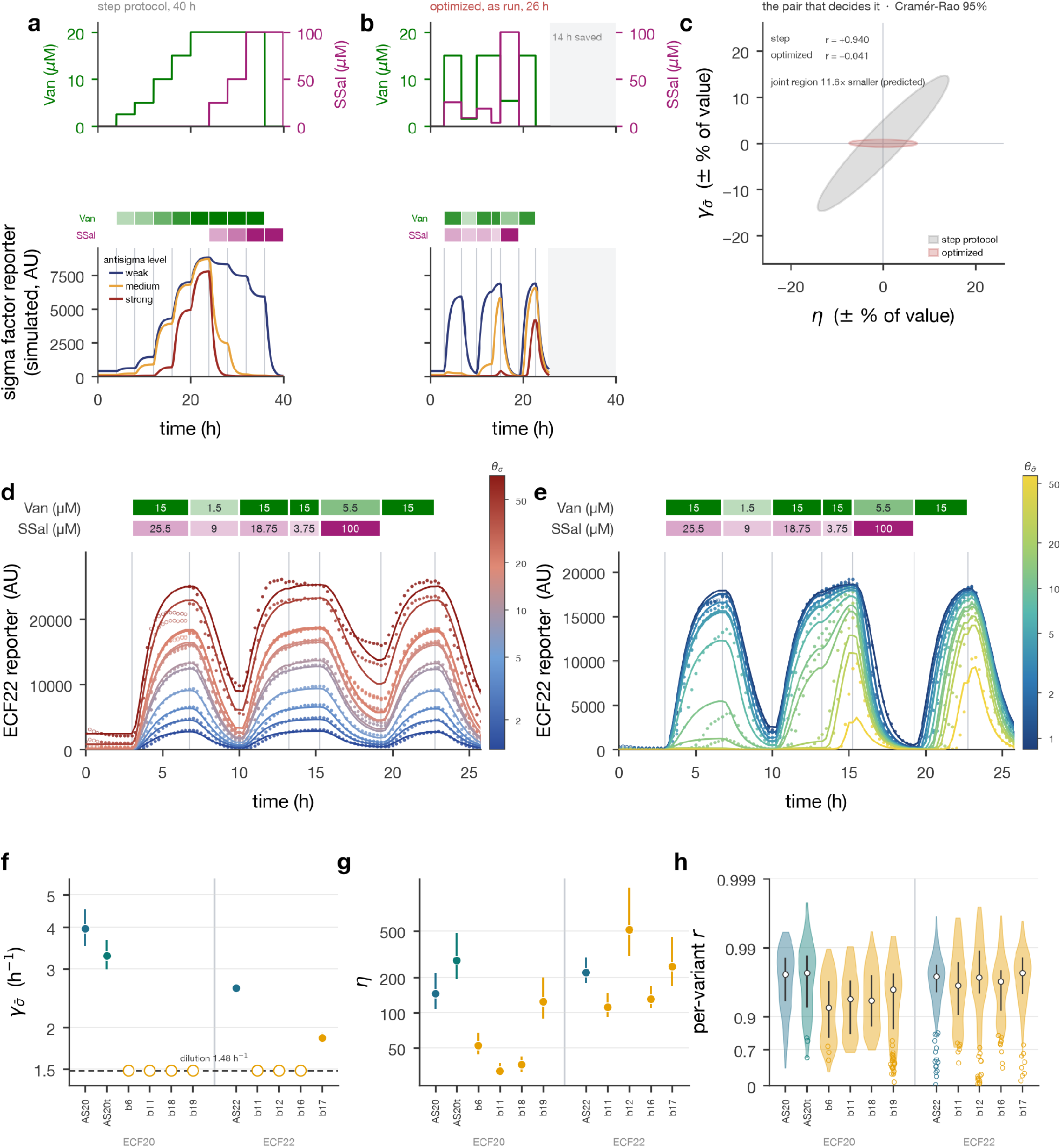
Optimized perturbations quantify expression, sequestration and turnover across binder architectures. **a,b**, Conventional 40-h step protocol (**a**) and model-optimized 26-h protocol used experimentally (**b**). Upper plots show inducer inputs; lower plots show simulated reporter responses at three antisigma expression levels. **c**, Predicted local Cramér–Rao 95% regions for the nominal ECF22 model, in log-parameter coordinates displayed as fractional uncertainty. The unrounded optimized schedule reduced correlation from 0.94 to *−*0.04 and joint area by 11.6-fold; these are design predictions, not empirical intervals. **d**, ECF22 reference trajectories across the *σ*-expression ladder in the presence of a non-cognate antisigma. **e**, ECF22-b12 trajectories across the antisigma-expression ladder at ECF22 RBS BCD-8.5k. Points are observations and lines are fits; open points were excluded from the loss. **f,g**, Antisigma removal (**f**) and effective sequestration (**g**) for 11 architectures. Whiskers delimit a 1% increase in loss when varying the indicated parameter with all others fixed, not confidence intervals. The dashed line is the imposed dilution reference; open symbols mark lower-bound removal estimates, for which *η* is conditional on that constraint. Sequestration rates are comparable within, not between, *σ* systems. **h**, Per-variant trajectory correlations for 1,453 variants above the detection limit. White circles and bars denote medians and interquartile ranges; open points show 80 below-detection variants excluded from the summaries.

To choose these perturbations, we used optimal experimental design ^53,54^. Using models fitted to the preceding experiments, we simulated candidate inducer schedules and searched for changes in concentration and timing that would make alternative parameter combinations produce distinguishable trajectories. The design considered several kinetic and reporter parameters together, while accounting for uncertainty in the expression ladders rather than treating expression levels as precisely known.

The resulting schedule varied both inputs non-monotonically over 26 h (Fig. 6b). Compared with the conventional 40-h step protocol (Fig. 6a), it was predicted to distinguish sequestration from removal more effectively despite requiring 14 h less experimental time. For the nominal ECF22 model, the unrounded optimized schedule reduced the predicted joint uncertainty region for *η* and 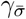 by 11.6-fold and largely removed their predicted correlation (Fig. 6c). These calculations describe the information expected under the preliminary model and assumed measurement noise, not empirical confidence intervals. The *D*_*s*_ design criterion, its parameter targets, and the rounding used for experimental implementation are described in the Supplementary computational methods.

We first estimated sigma production and basal activity across the expression ladder, together with the reporter production scale, using reference variants carrying a non-cognate antisigma (Fig. 6d). Reporter delay and inducer-response constants were inherited from the earlier calibration. With these reference parameters fixed, we jointly inferred each binder’s production ladder, basal activity, effective sequestration rate and removal rate. The optimized perturbation resolved responses across both expression ladders, including early differences that disappeared later as sigma production overwhelmed the binder (Fig. 6e; Supplementary Fig. S5). Across 1,453 variants above the detection limit, the median per-variant trajectory correlation was *r* = 0.965 (Fig. 6h). Saturation-affected measurements and the first hour were excluded from fitting; 80 low-signal variants were retained in fitting but excluded from the correlation summaries. Full-grid fits for all 11 architectures are provided in Supplementary Data 1, and fitted expression and basal-activity parameters in Supplementary Fig. S6.

The inferred removal rates were higher for the native-derived components than for most designed binders. AS20, AS20t and AS22 had fitted removal rates of 3.95, 3.28 and 2.62 h^−1^, respectively, compared with the imposed dilution-dominated reference of approximately 1.48 h^−1^ (Fig. 6f). Notably, truncating AS20 substantially reduced its measured stress response without returning its inferred removal rate to the dilution limit. By contrast, seven of eight de novo binders reached the dilution constraint, indicating that no additional removal was required to explain their trajectories. ECF22-b17 was the sole designed exception, with an inferred removal rate of 1.86 h^−1^.

Effective sequestration rates varied broadly among binders within each *σ* system (Fig. 6g). For the seven architectures whose removal rates reached the dilution constraint, however, the corresponding estimates of *η* are conditional on the assumed dilution rate because the two parameters remain coupled when that constraint is relaxed. Moreover, antisigma abundance was not calibrated on an absolute concentration scale. Values of *η* are therefore useful for comparisons among binders acting on the same *σ* factor, but not for absolute comparisons between the ECF20 and ECF22 systems. Despite these limitations, the measurements provide a kinetically diverse set of binders in which effective sequestration can be varied without the enhanced removal associated with the native components, supplying the parameterized parts used in the final closed-loop library.

Together with the earlier reporter calibration, the designed experiment supplied effective sequestration and removal rates, basal-expression estimates and binder-specific production ladders across all 11 architectures. The faster inferred removal of native antisigma factors was consistent with the proposed host-mediated turnover mechanism, although these measurements do not establish that stress activation caused their degradation. The experiment also provided the quantitative foundation for the final feedback library: calibrated production activities supplied the parameter axes for elasticity calculations, while independently inferred binder kinetics allowed us to investigate how component properties shaped closed-loop regulation and stability.

## 7 Mapping the design landscape of AIF circuits reveals robust adaptation, oscillatory instability, and nonlinear feedback attenuation

We next tested the hypothesis that interactions between native antisigma factors and the host contributed to the anomalous behavior of the first feedback library in Section 4. The stress responses and elevated inferred turnover found in the preceding experiments supported this explanation, but could have been incidental to the closed-loop defects. We therefore rebuilt the library with native, truncated and designed binders to test whether replacing the native inhibitors would reduce the anomalous disturbance responses while retaining effective feedback regulation. To do this, we assembled a combinatorial circuit library spanning 25,920 variants across five design dimensions.

The library combined 12 process-expression levels (*θ*_*p*_), 12 controller-expression levels (*θ*_*c*_), six disturbance-expression levels (*θ*_*d*_), six controller species (the four selected ECF20 binders together with native and truncated AS20) and five disturbance species (the four selected ECF22 binders together with native AS22). We recovered 22,448 unique variants, representing 86.6% of the designed library. Missing variants were concentrated in combinations of AS20 or AS20t with AS22 or ECF22-b17. The principal comparisons use ECF22-b16 as the disturbance binder, providing a common background with 96.6% coverage across the six controller architectures. Circuits containing this disturbance binder were also comparatively less prone to oscillation; sensitivity and oscillation maps across all five disturbance binders are provided in Supplementary Figs. S7 and S8. We write binder pairs as controller/disturbance, so ECF20-b19/ECF22-b16 denotes an ECF20-targeting controller binder paired with an ECF22-targeting disturbance binder.

We first tested how well the rebuilt circuits compensated for changes in the RBS controlling process expression. After correcting for reporter saturation, the independently measured noncognate open-loop reference responded approximately proportionally, with a median elasticity of 1.05. Feedback reduced this sensitivity, but to different degrees depending on the controller binder (Fig. 7a). ECF20-b19 gave the smallest process response, with a median elasticity of 0.13, while native AS20 and truncated AS20t also substantially attenuated the response. ECF20-b6 showed an intermediate response, whereas ECF20-b18 and ECF20-b11 retained greater sensitivity and were effectively tied. Signal filtering left different retained expression ranges, so these comparisons do not establish a ranking across the entire designed ladder. The designed binders therefore did not form a uniformly improved class: their reduced stress-reporter responses were compatible with both strong and relatively weak regulation.

**Figure 7.**
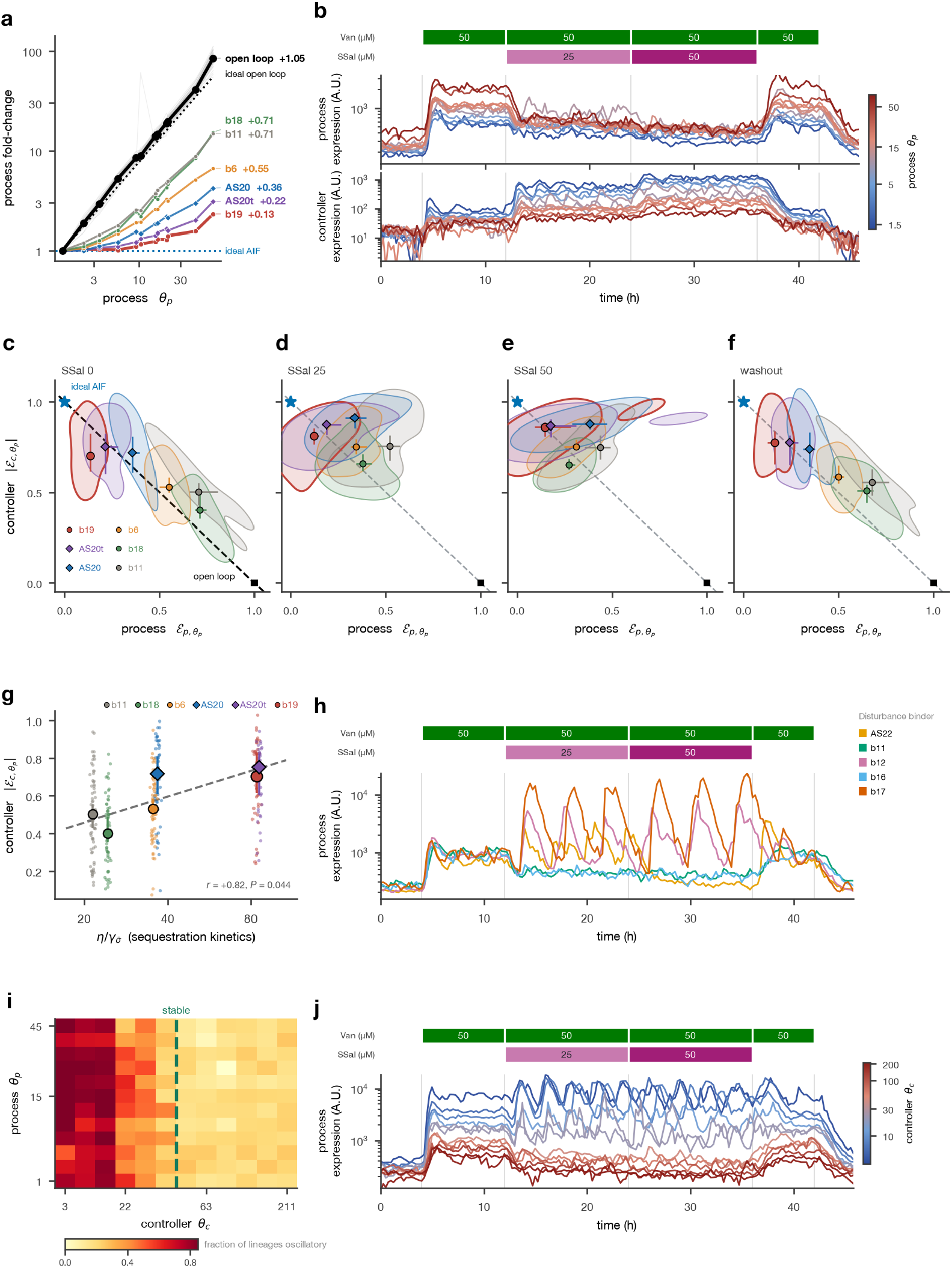
The feedback design landscape reveals disturbance-dependent compensation and sustained oscillations. **a**, Predisturbance process fold changes, normalized to the weakest retained activity within each fixed-background slice. Slopes are median slice-wise Theil–Sen estimates; the noncognate open-loop reference was measured independently. **b**, Variant-median process (mVenus) and controller (mJuniper) trajectories across *θ*_*p*_ for ECF20-b19/ECF22-b16, with *θ*_*c*_ and *θ*_*d*_ fixed. **c–f**, Paired elasticities before disturbance, at 25 and 50 *µ*M salicylate, and after withdrawal, respectively (324, 388, 388 and 316 slices). Contours enclose the densest 50% of estimates; the diagonal marks zero-disturbance complementarity. Slice membership can differ between phases. **g**, Predisturbance controller-elasticity magnitude versus independently fitted 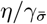. Small points are slices. Pearson correlation compares log kinetic ratio with linear elasticity (*r* = 0.82, *P* = 0.044, *n* = 6). Elasticities in **a,c–g** use channel-specific reporter desaturation (Supplementary computational methods). In **c–g**, symbols and whiskers show binder medians and 95% slice-bootstrap intervals; diamonds/circles denote native-derived/de novo binders. **h**, Process trajectories with different ECF22 disturbance binders and a fixed ECF20-b19 feedback core. **i**, Oscillation prevalence across *θ*_*p*_ *× θ*_*c*_ at 25 *µ*M salicylate. Cells show median per-variant fractions of oscillatory lineages across *θ*_*d*_; the dashed line marks the operational stability boundary. **j**, Process trajectories across *θ*_*c*_. Except **h** and the open-loop reference in **a**, panels use ECF22-b16.

The representative ECF20-b19/ECF22-b16 circuit showed a different disturbance response from the original example in Fig. 4g. Salicylate induction lowered process output rather than producing a sustained elevation above its predisturbance level, while controller activity increased (Fig. 7b). It also brought process outputs from different process-expression variants closer together. The disturbance therefore changed not only the output level, but also how effectively the circuit compensated for differences in process expression.

The paired sensitivity measurements showed that this effect extended beyond the representative circuit (Fig. 7c–f). These plots distinguish residual process sensitivity from the compensating controller response: ideal feedback occupies the upper-left corner, where process output is unchanged and the controller absorbs the change in process expression. Before disturbance, AS20t, AS20 and ECF20-b19 showed the largest controller responses, with overlapping bootstrap intervals. Across 324 expression slices, the median sum of process elasticity and controller-elasticity magnitude was 1.10, close to the small-disturbance complementarity benchmark of 1 (Supplementary Eq. S25). At 25 *µ*M salicylate, median controller compensation increased for all six binders. The largest reductions in process sensitivity occurred for ECF20-b6, ECF20-b18 and ECF20-b11, whereas ECF20-b19 and AS20t already showed relatively strong compensation before disturbance. After salicylate withdrawal, the binder groups generally returned toward their predisturbance positions. These are phase-level comparisons rather than paired equilibrium estimates: signal exclusions can change the contributing slices, and oscillatory variants remain included.

Independently measured binder kinetics were also associated with these differences in compensation. Across the six controller architectures, the normalized sequestration rate 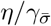 from the preceding open-loop experiment was positively associated with predisturbance controller-elasticity magnitude (Pearson *r* = 0.82, *P* = 0.044, *n* = 6; Fig. 7g). Thus, the kinetic characterization provided information about closed-loop behavior without refitting the components in feedback. With only six architectures, however, this association does not establish a quantitative prediction from the kinetic ratio alone; the concentrations and operating state of the circuit also matter.

The improvement after disturbance suggests that some circuits were initially producing too much controller inhibitor, rather than too little. The controller inhibitor (*c*_2_) is produced in response to process activity. If it accumulates, it suppresses the controller sigma factor (*c*_1_), reducing the controller’s ability to compensate. Crucially, sequestration need not eliminate controller activity completely. The fitted parameter *η* describes effective sequestration rather than directly measuring the molecular association rate. Rapid reversible binding, together with turnover of the bound complex, can also produce a finite effective sequestration rate. Our irreversible model is therefore a reduced description that does not require molecular binding itself to be slow or irreversible (Supplementary computational methods). Within this description, some free controller can remain even in the presence of excess inhibitor. A circuit can therefore produce a measurable output while operating in a state with attenuated feedback.

The full-model calculation shows why controller responsiveness depends nonlinearly on the operating state: sequestration depends on both controller and inhibitor abundance and competes with their independent removal. Too little inhibitor produces little compensatory action because sequestration contributes little to controller removal. With a large inhibitor pool, much of inhibitor production instead replaces molecules lost through dilution or degradation, weakening the shared production–sequestration balance required for near-ideal integration. When controller input and sequestration are sufficiently strong relative to independent removal, responsiveness has a maximum between these regimes (Supplementary Eq. S32). Increasing inhibitor production beyond this intermediate regime can therefore reduce responsiveness rather than strengthen it.

This provides a possible explanation for disturbance-dependent engagement of feedback. By suppressing the process, the disturbance also reduces production of the controller inhibitor. An initially inhibitor-loaded circuit can then move toward a more responsive operating state: the controller becomes more active, and differences in process expression are more effectively compensated. We describe this as windup-like because accumulated inhibitor attenuates controller responsiveness and reducing that accumulation can restore it. The mechanism does not require a separately saturating actuator, as in conventional windup analyses ^55^, and the present measurements do not directly establish the inhibitor pool or a prolonged recovery time. Rather, the calculation supplies a mechanistic interpretation of why a disturbance could strengthen regulation instead of simply exhausting its capacity.

Even circuits built with designed binders could switch from relatively steady output to large-amplitude, sustained oscillations after disturbance induction (Fig. 7h). In circuits with ECF20-b19 as the controller binder, changing only the ECF22 disturbance binder while keeping the process and controller components and their expression settings fixed produced responses ranging from relatively steady regulation to oscillation. The onset therefore depended on the disturbance branch, not solely on the feedback core. These responses provide an experimental counterpart to the theoretical prediction that antithetic feedback can destabilize the process it regulates ^22,37^. Matched process-expression sweeps across all binder pairs show the wider range of responses (Supplementary Fig. S9).

Quantifying oscillations in individual lineages revealed which expression settings made circuits susceptible to this disturbance-induced transition. In the ECF20-b19/ECF22-b16 series, oscillation prevalence depended much more strongly on controller expression than on process expression (Fig. 7i). Low controller expression supported frequent oscillations under disturbance, whereas higher expression suppressed them, a transition also apparent in the direct controller-expression sweep (Fig. 7j). Before disturbance, every controller-expression level in this series remained below the operational oscillation cutoff. Across disturbance binders, the transition boundaries spanned more than a threefold range of controller activity at 25 *µ*M salicylate (Supplementary Fig. S8), showing that the disturbance branch shaped the expression range over which instability emerged.

The suppression of oscillations at higher controller expression is particularly informative because it contrasts with a familiar prediction from the rapid-sequestration limit. For an ideal antithetic controller coupled to a multistage linear process, this limit imposes an upper production-gain bound for stability. When sequestration is slower, however, the relationship can reverse, allowing increased production gain to stabilize the circuit ^37^. The observed transition follows this latter direction, although its much stronger dependence on controller than process expression indicates that production gain alone is insufficient to explain it. Independent controller removal and the dynamical disturbance branch also distinguish the present circuit from that theoretical model. The earlier analysis therefore provides a precedent for the direction of the transition, not a quantitative prediction of its location. Likewise, our controller-response calculation explains how disturbance could strengthen feedback, but does not determine whether the resulting response will be stable.

Together, these results support the rationale for replacing native antisigma factors: designed binders could retain strong compensation while reducing the associated stress responses and excess turnover. They do not establish that host interactions explain every anomaly in the first library. The redesigned circuits nevertheless revealed a landscape containing weak compensation, strong regulation and sustained oscillation, with disturbance changing both the output and the effectiveness of feedback.

This landscape shows that regulatory performance depends on molecular implementation and operating conditions, rather than being a fixed property of the feedback architecture. By relating systematic variation of circuit components to independently measured kinetics and dynamic responses, we could test where the behavior predicted by theory was realized and investigate the departures. The hierarchy thus connected component characterization to a map of circuit function and its limits, providing a basis for designing and refining feedback systems across a range of conditions.

## 8 Discussion

A central challenge in biological engineering is translating theoretical predictions into experiments that can test and refine our understanding of how living systems work. A model can make precise predictions about how a circuit should respond to a change in one of its parameters, but identifying a corresponding genetic perturbation and measuring its effect can require substantial experimental development. This practical separation between theoretical predictions and the experiments needed to evaluate them has been described as a phenomenological gap ^56^. BARRACUDA was developed to make this exchange between theory and experiment more accessible. We used antithetic integral feedback as a setting in which theoretical questions could guide the development of an experimental system, and the resulting measurements could, in turn, expose where the theory needed to be refined.

The elasticity calculation provides a concrete example of what this requires. Theory predicts how changes in process expression should be absorbed by a compensating controller response. To test that prediction, however, we need to know how much expression was changed. Replacing an RBS gives us a different circuit, but its sequence alone does not supply the denominator in a sensitivity calculation. The preceding open-loop measurements established that denominator through calibrated RBS activities, allowing us to compare the measured responses of the process and controller with theoretical expectations. Characterizing the simpler systems therefore made it possible to ask a quantitative question about the more complex ones. This dependence on earlier measurements recurred throughout the study, culminating in binder kinetics that were measured in open loop and then associated with differences in closed-loop regulation, without any closed-loop refitting.

The experimental design also made it possible for observations to change the direction of the work. Following individual lineages through repeated perturbations revealed oscillations that were obscured by population aggregation, while structured variation across the libraries showed how these dynamics depended on circuit design. Together with sustained overadaptation, these observations exposed limitations of the initial model and motivated experiments on host stress and binder turnover. Theory then helped determine how to perform those experiments. Model-designed input sequences separated kinetic effects that were difficult to distinguish under conventional titrations, providing a basis for interpreting the redesigned components. The interplay between theory and experiment thus shaped both the questions we asked and the measurements available to answer them.

For biological control, these results emphasize how strongly the behavior of a feedback architecture depends on its molecular implementation and operating conditions. Independent removal of the controller species changes the balance required for integration ^37,40^, making cellular processing of a binder relevant alongside its ability to sequester its target. The final libraries allowed these properties to be compared across regimes of regulation and oscillation. Analysis of the competition between sequestration and independent controller removal further suggested how a disturbance could improve regulation by moving the controller into a more responsive regime. This explanation remains to be tested through direct measurements of controller-inhibitor accumulation, and quantitative accounts of the oscillatory transitions and of the sustained overadaptation seen in the first feedback library are both still missing. These are specific problems for further theory and experiment that emerge from the same framework.

The scope of the resulting models is necessarily tied to the measurements. The whole-library residuals illustrate that a shared expression model captures broad trends without removing all component-context dependence (Supplementary Fig. S2). Our calibrations describe effective activities in a particular cellular context, and fluorescence does not establish an absolute molecular concentration scale. Transferring these circuits to another strain or growth condition will require renewed characterization. Even within the present study, some component properties transferred more faithfully than others. A useful theory of biological engineering will need to accommodate this context dependence, with experiments establishing which descriptions remain predictive as the system changes.

We view synthetic feedback circuits in *E. coli* as a tractable setting in which to develop this way of working. The broader opportunity is to make sustained interaction between theory and experiment practical across a growing range of biological engineering problems. A theoretical prediction becomes more useful when the experiment needed to test it can be carried out, and an unexpected observation becomes more informative when we can systematically vary the conditions that produced it. By bringing these activities into a common experimental framework, BARRACUDA provides a basis for circuit design in which each round of construction and measurement contributes to a more general understanding of the system.

## Materials and Methods

### Bacterial strains and growth conditions

Experimental strains were derived from *E. coli* MG1655 through the parental strain LIB591. This strain combines the Marionette WT regulatory cassette ^32^, the CymR–T7 RNA polymerase integration used for barcode transcription in MARLIN ^19^, and three chromosomal landing pads ^57^. Library-specific chromosomal constructs, including fluorescent markers and sigma-dependent reporters, were introduced through the kanamycin-associated landing pad using an integration workflow adapted from clonetegration. The Marionette and CymR–T7 elements were retained across experimental hosts. Chromosomal mKate2Hyb provided the constitutive segmentation marker, while yellow- and/or cyan-channel reporters measured circuit activity. The designation “mCherry” in acquisition configurations and some analysis outputs denotes the optical channel, not the fluorescent protein expressed by these strains. Experiments were performed at 37 °C in MOPS EZ Rich Defined Medium with glucose (Teknova, M2105). Circuit plasmids carried chloramphenicol resistance.

### Combinatorial assembly and barcode insertion

Libraries were assembled through an intermediate, unbarcoded circuit library. Promoter, translation-initiation, coding-sequence and terminator parts were manually pooled at approximately equimolar input ratios for each variable position. Transcriptional units were assembled using BsaI, amplified and gel-purified, then combined into circuit backbones using BbsI. The unbarcoded library was recovered in liquid culture before barcode insertion. This intermediate checkpoint separated assembly of the circuit from assignment of its optical barcode and allowed the circuit library to be recovered independently of the final experimental host.

In the modified backbone, the barcode placeholder carried NahR, VanR and LacI. These temporary regulators enabled inducible circuit libraries to be propagated in commercial NEB 10-beta electrocompetent cells before transfer into the experimental host. The placeholder was subsequently excised using BsaI, the full-length backbone was gel-purified, and MARLIN barcode cassettes were inserted by a two-part Golden Gate assembly. This step removed the temporary regulatory cassette. Barcoded assemblies were transformed into the experimental strain or the Marionette cloning strain. The final barcode construct retained both the original MARLIN terminator and an additional T7-specific terminator.

Transformant yields were estimated by plating a 10^−3^ dilution of the transformation recovery. For the initial unbarcoded assembly, the undiluted recovery was used to establish the liquid library, with a target yield of at least ten transformants per theoretical circuit variant. For barcode insertion, several dilutions of the recovery were used to establish bottlenecked liquid cultures targeting approximately 10–20 barcoded transformants per theoretical variant. This limited barcode complexity while providing repeated sampling of circuit variants. These ratios describe assembly targets, not verified uniform representation or guaranteed coverage; subsequent sequencing established the recovered barcode–genotype associations.

### Microfluidic culture and programmable inputs

Mother-machine devices, cell loading and on-chip culture followed the MARLIN workflow ^19^. Cells were retained in growth trenches supplied from feeding channels, allowing lineages to be followed during programmed changes in medium composition. Environmental inputs were controlled by an Arduino-driven set of Precigenome solenoid valves and Longer peristaltic pumps. Each supply line was set to approximately 15 mL h^−1^, with only one input valve open at a time. Pulse-width modulation over a 60-s period varied the fraction of time assigned to each input while maintaining the same nominal flow rate into the chip. The duty fraction of each input therefore determined its contribution to the time-averaged medium composition. Successive input streams mixed along the tubing and at the shared inlet rather than in an actively stirred chamber.

Mixing was checked by supplying fluorescein under different switching conditions and measuring fluorescence in the feeding channel. Switching periods of 60 s or less produced consistent feeding-channel fluorescence without detectable periodic intensity oscillations under the tested conditions. This measurement assessed smoothing of the switched inputs; it was not a calibration of the absolute concentration or transport delay of each inducer. Vanillic acid, sodium salicylate and IPTG schedules were specified separately for each experiment, as indicated in the corresponding figures.

### Time-lapse fluorescence microscopy

Imaging used the Ti7 microscope setup, comprising a Nikon Ti2-E body, a Nikon 20 × 0.8 NA Plan Apo Lambda D objective, a Hamamatsu ORCA-Quest 2 camera and a Lumencor Spectra III light source. Images were acquired at 15-min intervals with 40-ms exposures in each fluorescence channel. Illumination intensity was selected separately for each experiment. The constitutive mKate2Hyb signal was used for segmentation, and yellow- and cyan-channel signals were used for phenotyping as required by the library. Experiment-specific intensity settings therefore did not define a common absolute fluorescence scale across datasets; the normalization and cross-experiment calibration used in individual analyses are described in the Supplementary computational methods.

### Optical barcode decoding

Libraries used the 30-bit barcode architecture and sequential fluorescence in situ hybridization workflow described for MARLIN ^19^. Following phenotypic imaging, T7-driven barcode transcripts were decoded in the device to associate imaged lineages with their circuit identities through the sequencing-derived codebook. The barcode architecture was unchanged apart from the additional T7 terminator described above. Modifications to the liquid-handler protocol included adjustments to step timing, additional washes and simplified buffer formulations.

### Long-read sequencing and barcode–genotype assignment

Circuit–barcode associations were determined using Oxford Nanopore PromethION sequencing. The Harvard Bauer sequencing core supplied basecalled FASTQ files; local analysis began from these reads and did not include basecalling or duplex-read generation. Where sequencing runs contained multiple libraries, reads were separated by library before downstream processing.

Reads were processed with the Nextflow-based `barcoded_amplicon pipeline` (https://github.com/shenker/barcoded_amplicon_pipeline). The workflow uses graph references to represent the barcode and combinatorial circuit sequences, groups reads by barcode alignment, constructs consensus sequences and aligns those consensuses to the circuit reference to identify component combinations. This replaced the sequencing-processing implementation used in MARLIN while retaining the barcode architecture. Graph paths provided both the circuit assignment and its 30-bit barcode for export to the optical-decoding codebook.

For the archived 3 April 2026 pLIB578 run, the saved configuration specifies abPOA consensus generation, with contributing reads limited to 10,000 bases and a maximum alignment-divergence setting of 0.3; the execution log records Nextflow 24.04.4. The codebook-export script retained primary alignments spanning the designated amplicon endpoints and read groups with depths of 15–100. These settings describe that run and are not assumed to apply to every library.

### Image processing and quantitative analysis

Image processing and optical decoding used the MARLIN workflow ^19^, with project-specific software changes. Segmentation based on the chromosomal mKate2Hyb marker supported extraction of lineage-resolved fluorescence trajectories, which were linked to circuit genotypes through the barcode codebook. Individual lineage measurements were retained for analyses of temporal variability and oscillation prevalence. Variant- and phase-level summaries were used where required for calibration and model fitting; their definitions and inclusion criteria are given with each analysis in the Supplementary computational methods.

The local-background correction, low-signal handling, fluorescence-response models, parameter inference, optimal experiment design and experimental elasticity calculations are specified in the Supplementary computational methods. In particular, the constitutive segmentation marker was not treated as interchangeable with the background-corrected yellow and cyan reporters.

### Experimental replication and uncertainty

Each final library dataset was obtained from a single pooled microfluidic imaging experiment, with variants measured concurrently under a shared input schedule. Within-run replication comprised multiple lineages carrying the same circuit variant. Independent repeat imaging experiments were not performed for each library. The pooled design permits comparisons within a common experimental environment, but does not quantify between-experiment variability. Resampling units and uncertainty calculations differ by analysis and are specified in the Supplementary computational methods; repeated time points within a lineage were not interpreted as independent experimental runs.

## Author contributions

N.O. and J.Q.S. initiated the project, with conceptual input from J.P. J.Q.S. developed the sequencing pipeline for assigning circuit genotypes to barcodes and generating the codebook used for FISH decoding. N.O. developed the experimental platform and combinatorial libraries, performed all cloning and experiments, developed the remaining project-specific software, analyzed the data, and prepared the figures. D.C. provided the stress-response promoter sequences and constructs. N.O. and J.P. wrote the manuscript.

## Acknowledgements

The authors would like to acknowledge Carlos Sanchez, Daniel Eaton, and Niko McCarty for their helpful feedback on the manuscript. We would also like to thank Yonatan Chemla and Jai Padmakuma from Chris Voigt’s lab for sharing strains and cloning protocols with us.

## Supplementary computational methods and theory

### S1 Scope and parameter conventions

The computational methods follow the experimental progression from constitutive and inducible expression calibration (Fig. 2), through sigma–antisigma characterization (Fig. 3) and binder screening (Fig. 5), to optimal experiment design and dynamic kinetic inference (Fig. 6). The feedback-theory sections then derive the predictions used to interpret the closed-loop libraries. Finally, we describe fluorescence-background correction, experimental elasticities and lineage-resolved oscillation analysis (Figs. 4 and 7). Each analysis has its own parameter-sharing structure, residual definition and uncertainty calculation, as specified below. Symbols and their molecular meanings are collected in Supplementary Table S1.

Expression activities are denoted by *θ* in the manuscript. The analysis files use `beta` for the inducible reporter production ladder and `rho`/`rho as` for sigma/antisigma production ladders. The fluorescent protein stored as `rcVenus` is displayed as mVenus. RBS identifiers beginning with `SD2` are labels for the BCD series used here; they are not numerical estimates of expression activity.

### S2 Constitutive expression calibration

For promoter *i*, RBS *j*, and reporter *k*, fluorescence was modeled as

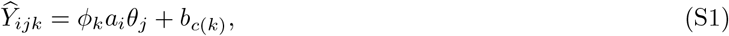

where *b*_*c*(*k*)_ is shared by reporters measured in the same imaging channel. The figure builder selects the 12-member BCD series from the constitutive library and fits the four reporters jointly. The inputs are the channel-specific, normalized construct measurements supplied by the static measurement table.

The residual for each observation is *r*_*ijk*_ = log *Y*_*ijk*_ − log *Ŷ*_*ijk*_. The implementation minimizes the `soft l1` robust objective applied to these residuals, using `scipy.optimize.least_squares` ^58^, a lower parameter bound of 10^−6^, and a maximum of 2,000 function evaluations. The two multiplicative scale freedoms in Eq. S1 are removed by fixing one promoter activity and one RBS activity to unity during fitting. These parameters are optimized on their positive, bounded natural scale; taking logarithms of the residual does not imply that the optimization coordinates are logarithmic. Goodness of fit is calculated in log fluorescence space.

Only positive measurements below the within-reporter 99th percentile enter the fit, retaining 555 of 563 measured BCD-library constructs and giving *R*^2^ = 0.9238. Supplementary Figs. S1 and S2 use the same model and filtering procedure applied jointly to all 26 RBSs, retaining 1,172 constructs (*R*^2^ = 0.913). The whole-library fit re-estimates all shared activities; its 324-fold range for the Jansen BCD family should not be substituted for the 392-fold range from the BCD-only fit in Fig. 2.

### S3 Inducible reporter calibration

Each reporter was fitted independently across pVanCC, pSalTTC and pTac. The unit-amplitude input response

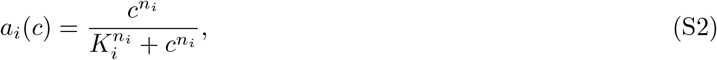

where *c* is inducer concentration, *K*_*i*_ the half-maximal concentration and *n*_*i*_ the Hill coefficient. The notation *H*_*i*_(*I*) below is equivalent to *a*_*i*_(*c*), with *I* denoting inducer concentration. Basal activity is additive and outside this response. The Hill amplitude is fixed to unity, with the production scale carried by *θ*_*j*_; sharing *θ*_*j*_ across promoters imposes a common inducible production increment for a given RBS within each reporter.

For promoter *i* and RBS *j*, the dynamics are

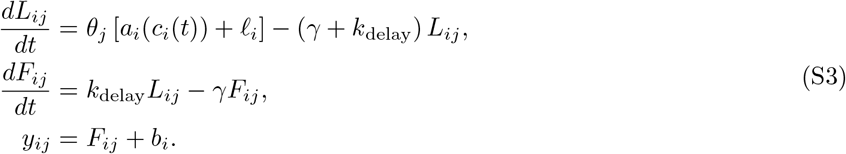

Here, *L*_*ij*_ is a latent reporter state, *F*_*ij*_ the observable fluorescent state, *l*_*i*_ the promoter’s basal activity and *b*_*i*_ its fitted background. The loss rate is *γ* and the transition rate is *k*_delay_. The delay is phenomenological and is not a direct measurement of fluorophore maturation. There is no additional sigma state or sum of production rates across RBS variants in an individual trajectory.

Within a reporter, *γ* and *k*_delay_ are shared across trajectories; *K*_*i*_, *n*_*i*_, *l*_*i*_, *b*_*i*_ are promoter-specific; and *θ*_*j*_ is shared across promoters at a given RBS. All parameters are fitted independently for mVenus and mJuniper because of coding-sequence-dependent expression. Initial conditions are the uninduced steady state, *L*_*ij*_(0) = *θ*_*j*_*l*_*i*_*/*(*γ* + *k*_delay_) and *F*_*ij*_(0) = *k*_delay_*L*_*ij*_(0)*/γ*.

The data adapter subtracts a channel-specific second-percentile background at each timepoint and clips negative corrected values to zero. It masks a trajectory from the first raw observation at or above 60,000 fluorescence units onward. The weakest BCD element, BCD-0.1k, showed no detectable mJuniper induction across the tested promoter and inducer conditions and was excluded from dynamic fitting, leaving 11 mJuniper and 12 mVenus RBS activities. This exclusion was specific to the RBS–reporter combination; BCD-0.1k was not removed from other expression contexts solely on this basis.

Trajectory residuals are

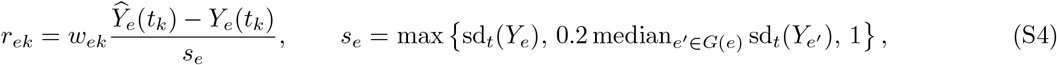

where *G*(*e*) comprises trajectories with the same indexed RBS variant and *w*_*ek*_ is zero for masked observations and one otherwise. The scale is derived from trajectory variation; it is not an independently measured standard error of the observation.

The per-reporter fit is initialized from independently fitted promoter models and refined by trajectory matching. The inspected implementation uses a trust-region reflective least-squares optimizer ^59^ in log-parameter coordinates, a sparse finite-difference Jacobian, and Radau integration ^58^ with relative and absolute tolerances 10^−6^ and 10^−8^. The saved results used by the figure are `inference_results_real_1b_per_fp`; their stored loss rates are 1.492197 and 1.481595 h^−1^ for mJuniper and mVenus, respectively. Their effective delay rates are 6.932638 and 2.785284 h^−1^.

For dose-response visualization, the dynamically inferred *K*_*i*_, *n*_*i*_ are retained and two affine coefficients are estimated for each variant against *H*_*i*_(*I*). Normalized plateau values are obtained using these coefficients. This visualization assesses a shared response shape and is not an independent parameter-free prediction.

### S4 Joint steady-state inference for the sigma–antisigma screen

The general dynamics used to describe an isolated regulatory pair are

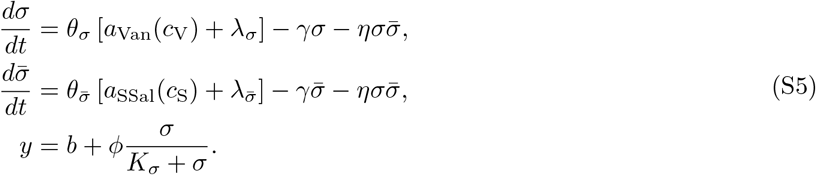

Here, *a*_*i*_(*c*) is the unit-amplitude response in Eq. S2. The inducer-response constants are inherited from the reporter screen, whereas basal activities are refitted in each regulatory context. The *λ* terms take the place of the reporter-context *l*_*i*_ and are not added to inherited basal activities. Effective RBS activities are *θ*_*σ*_ and 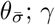 is dilution-dominated loss and *η* is an effective sequestration rate represented by irreversible mutual removal in this reduced model. It is not a direct measurement of the molecular association rate. The reporter has output scale *ϕ*, background *b* and half-maximal free-sigma concentration *K*_*σ*_.

Protein identities and RBS indices are suppressed in this dynamical notation. In fitting, sigma basal activity is RBS-specific, *λ*_*σ,i*_, while antisigma basal activity is shared across its ladder. Fixing *K*_*σ*_ = 1 defines a system-specific concentration scale; effective activities support quantitative fold-range comparisons within a pairing, not absolute concentration comparisons across different sigma reporters.

The screen in Fig. 3 is fitted to plateau measurements using the algebraic steady state of the sequestration model. The fitting routine does not integrate an ODE for each optimization evaluation. The plateau extractor takes the median of the last third of unmasked samples within each phase, retaining at least three samples when available and omitting phases with fewer than three unmasked samples. Let

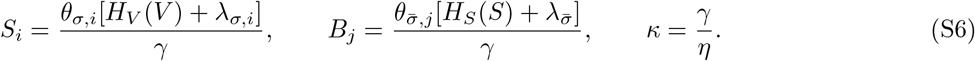

Then the free sigma concentration satisfies

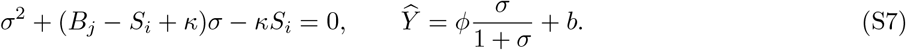

Writing *w* = *B*_*j*_ − *S*_*i*_ + *κ* and *d* = (*w*^2^ + 4*κS*_*i*_)^1*/*2^, the positive root is evaluated as 2*κS*_*i*_*/*(*w* + *d*) when *w* ≥ 0 and (*d* − *w*)*/*2 otherwise, avoiding subtractive cancellation. The parameter called `K_d` in this implementation is *κ* = *γ/η*, derived from the reduced sequestration model at steady state. Its relation to an equilibrium dissociation constant requires assumptions about complex turnover; the parameter name alone does not establish a microscopic binding interpretation. The relation to reversible binding is discussed in Supplementary Section S9. The input Hill functions are fixed from the mJuniper calibration: (*K*_*V*_, *n*_*V*_) = (34.24 *µ*M, 2.178) and (*K*_*S*_, *n*_*S*_) = (59.28 *µ*M, 1.695). The loss rate is fixed to *γ* = log(2)*/*(28*/*60) h^−1^, the dilution rate for a 28-minute doubling time and consistent with the reporter loss rates recovered independently in Fig. 2h, and *K*_*σ*_ = 1 fixes the concentration scale. For *n* sigma and *m* antisigma rungs, each pair has 2*n* + *m* + 4 free parameters: 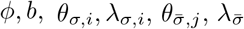, and *κ*.

Reference and pair-specific observations enter a single residual vector. For cognate pairs, the reference arm pools certified noninteracting partners; for noncognate pairs, the tested partner is excluded from the reference pool. For observation *k* in arm *a*, the residual is

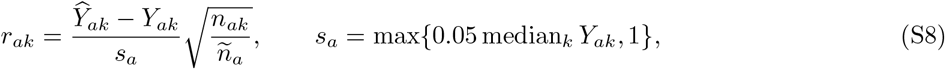

where *n*_*ak*_ is the associated trench count and *ñ*_*a*_ is the median count within that arm. No additional minimum-trench threshold is imposed in this fitting routine.

Pre-fit exclusions identify variants with weak induction inconsistent with their expression ladder, antisigma variants with markedly reduced suppression relative to weaker labeled rungs, and antisigma rungs with fewer than three surviving genotype combinations. The weakest surviving sigma rung is also removed iteratively if its pooled reference induction is less than twofold. Specifically, the sigma-variant rule requires both less than twofold induction and an induced level below one third of the brightest weaker sigma rung at the same antisigma rung. The antisigma rule requires suppression to be more than one decade weaker than a weaker antisigma rung that suppresses by more than half a decade.

Fitting uses bounded trust-region reflective least squares ^59^ in log-parameter coordinates with Jacobian-based coordinate scaling. One initial point is selected from a 20-point grid of 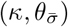 seeds, followed by 40 perturbed starts with the *κ* coordinate distributed across its log bounds. The saved summary records all 41 starts within 1% of the best cost for each of 25 pairs, with no recorded evaluation-limit failures or exceptions.

Asymptotic intervals are calculated from the weighted-residual Jacobian *J* with respect to log parameters. Coordinates within 10^−4^ of their log-box span from either boundary are excluded from the covariance calculation. For the remaining coordinates,

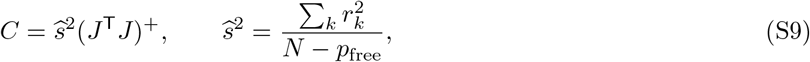

where + denotes the pseudoinverse. A parameter interval is 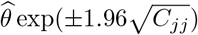. Displayed rungs require a 95% multiplicative half-width no larger than twofold and are rejected if their observed response never reaches twofold. For a ratio of the retained maximum and minimum activities, the variance of its logarithm is *C*_*hh*_ + *C*_*ll*_ − 2*C*_*hl*_. This interval conditions on the selected extreme rungs; it does not include uncertainty about which rung is the maximum or minimum. Ranges based on an incomplete retained ladder are reported as conservative bounds.

### S5 Computational analysis of the binder screen

#### Expression axes and reporter trajectories

The screen in Fig. 5 samples six binder RBS sequences from the calibrated ladder: BCD-0.2k, -0.75k, -2.3k, -8.5k, -18k, and -25k. Cross-talk is assessed using the other ECF reporter: ECF22 for binders targeting ECF20, and ECF20 for binders targeting ECF22. RBS sequences are mapped to the reference activities *θ*_*j*_ from the joint constitutive-expression calibration in Fig. 2, exported as `rbs strength.csv`. The same six-rung axis is used for specificity, constitutive-reporter burden and stress elasticities, for both targets and every native, truncated and de novo binder. All six screened sequences have a usable reference activity; none is excluded for lack of calibration. This part-level axis is an expression proxy, not a measurement of binder abundance or a binder-specific production rate, and does not depend on the later kinetic fits. Its use assumes that relative RBS activities transfer to the binder-expression context.

For the displayed reporter trajectories, a separate background estimate is obtained for each sigma-reporter system from the initial uninduced window and the terminal washout. Within each window, temporal medians are calculated per variant and then a median is taken across the selected control variants. The extraction selects controls whose reporter sigma differs from the binder’s designated target. Background is linearly interpolated between the two window midpoints and held at the nearest endpoint outside them. The subtraction is clipped at zero. This screen-specific correction is separate from the local image background correction used for the closed-loop experiments.

#### Specificity statistic

At each timepoint, the off-target reporter is pooled within each calibrated binder-expression rung using trench-count weights. Inputs with fewer than 20 trenches are excluded. Ordinary least squares fits log_10_(max{*Y*, 1}) against log_10_ *θ*_*j*_ when at least four rungs span at least 0.5 decades. The per-binder suppression statistic uses the mean slope over 13–20 h, with negative slopes indicating suppression of the noncognate reporter. The saved gate standardizes its negative by a pre-induction reference width of 0.0116, estimated using the same constitutive-expression reference axis.

An additional statistic detects separation of the RBS trajectories, including changes that do not produce a negative slope. Each rung’s background-subtracted reporter is divided by its own median over 5–11 h. The standard deviation across normalized rungs is computed at each timepoint and averaged over a window. The difference between 12–20 h and 5–11 h is divided by the population standard deviation of the pre-induction spread across the de novo binder library for that target. At least four calibrated rungs are required. Writing these standardized slope and spread statistics as *Z*_slope_ and *Z*_spread_, the specificity score is

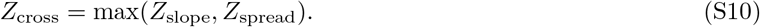

Binders with *Z*_cross_ ≥ 3 fail the specificity gate. The saved tables retain 16 of 20 ECF20 binders and 10 of 20 ECF22 binders. This cutoff is an operational screening rule, rather than a multiple-comparison-adjusted significance test.

The red trajectory annotations in Fig. 5c,d use a separate graded-suppression diagnostic. It flags a negative noncognate slope exceeding three reference standard deviations, or one exceeding 1.5 reference standard deviations accompanied by a monotone decline across the ladder (Spearman correlation at most −0.90 and a decrease of at least 0.08 in the normalized off-target reporter response). For this diagnostic, each rung’s response is its minimum over 13–20 h divided by its mean over 10–12 h; the decrease is measured from the weakest to the strongest RBS. This diagnostic can flag a shallow graded response that passes the composite gate; conversely, the composite gate can fail on rung separation alone. ECF22-b8 and ECF22-b6 illustrate these two cases, respectively. These annotations do not determine the reported specificity-pass counts. Supplementary Fig. S3a,b instead marks failures of the composite gate directly.

#### Constitutive-reporter burden

Burden is measured from raw constitutive mKate2Hyb in the cognate sigma background. The calculation again uses a log–log slope across calibrated RBS activities and a spread across trajectories normalized to their own pre-induction levels. For the slope, the baseline window is 0–4 h; for spread, it is 5–11 h. The larger signed slope and the larger spread from the 12–20 h and 16–21 h windows are retained, and the corresponding baseline statistic is subtracted.

The implemented burden standardization uses the distribution of these changes across de novo binders. For either change statistic Δ_*b*_,

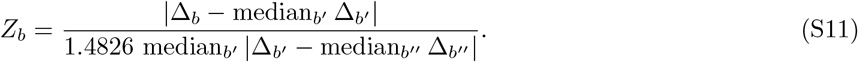

The scale factor 1.4826 makes the median absolute deviation a consistent estimator of the standard deviation under normality ^60^. The reference antisigma controls are excluded from the centering and scale calculation. The larger of the slope and spread scores is compared with 3. Among specificity-passing binders, this excludes ECF20-b12 and ECF22-b18. The burden score is thus standardized against library-wide changes, whereas the specificity score uses pre-induction reference widths.

#### Native and truncated reference controls

The four native and truncated controls are evaluated using the same estimators and normalizations as the de novo binders and are plotted for comparison; they are not included in the 20-design pass counts. The mean noncognate log–log slopes are +0.060 for AS20 and −0.029 for AS22, indicating reporter increases and suppression, respectively. Their cross-talk scores are 11.58 and 5.86, both determined by the rung-separation statistic rather than the slope statistic (*Z*_slope_ = −5.20 and 2.50, respectively). Their burden scores are 16.98 and 5.75. Both native controls therefore exceed both operational cutoffs, but the direction of their reporter responses differs. Neither the composite cross-talk score nor the constitutive-reporter burden proxy identifies the molecular mechanism or establishes direct cross-binding.

#### Stress-reporter elasticities

For each stress promoter, binder, and timepoint, the stress response is summarized by the slope of log reporter intensity against log calibrated RBS reference activity, denoted 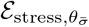. Here, 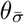 identifies the RBS driving binder expression: its numerical value is the reference activity *θ*_*j*_ from the joint constitutive-expression calibration (Fig. 2), exported as `rbs_strength.csv`. A single activity per RBS is shared across fluorescent reporters and promoters in that calibration. The same six calibrated rungs are used for all 44 binder genotypes and both targets, including native and truncated antisigma controls. No rung is omitted for lack of calibration, and no binder-specific parameter or surrogate ladder from the later kinetic fits enters the stress analysis. This shared part-level axis is an expression proxy, not a measurement of binder abundance or its protein-specific production rate. Specificity and burden slopes use this same reference axis, with their own response statistics and standardizations as described above.

A constant rescaling of the reference ladder changes the logarithmic intercept but not its slope. Relative rung spacing, however, is inferred from the constitutive-expression model; its uncertainty and transfer to the binder-expression context are not propagated into these elasticity estimates. At least four finite observations are required for the slope; stress intensity is clipped at 1 count before taking logarithms. Both axes use base-10 logarithms, giving a dimensionless slope equivalent to using natural logarithms on both axes. A centered five-timepoint running median smooths the slope series. The extraction retains all nine stress-promoter classes, including weakly expressing *P*_bamA_; there is no promoter-level brightness exclusion or absolute-intensity row filter. It checks individual variant labels against their pre-induction stress intensity: a variant is excluded from the stress analysis when another promoter’s median baseline is closer in natural-log intensity, the discrepancy from its assigned class exceeds 0.7, and the improvement from switching classes also exceeds 0.7. This flags a baseline inconsistency; it does not by itself prove a barcode assignment error. Native stress responses are reported as phenotypes and are not used as the constitutive-mKate2Hyb burden gate.

The operational reference threshold is recomputed from the current, smoothed elasticity series during 5–11 h, before salicylate addition. Pooling all 44 binder genotypes across both targets and their stress reporters gives 9,550 finite values, with population standard deviation 0.03918; three times the unrounded value gives the threshold *T* = 0.11754. The full-library reference is retained consistently for the standardized stress scores and integrated responses. This threshold is an operational reference, not a calibrated false-positive probability. Timepoints are correlated, the series are smoothed, and basal expression can contribute before induction. The gray bands in Fig. 5e and Supplementary Fig. S3c are drawn at this same ±*T* threshold, which is computed once and shared by the elasticity panels and the integrated scores.

For Supplementary Fig. S4, panel a takes the maximum signed elasticity over the recorded time course and over the promoters in each regulon. Panel b restricts scoring to 12 h onward, after salicylate addition, and integrates the positive excess on the promoter carrying each binder’s largest post-induction peak:

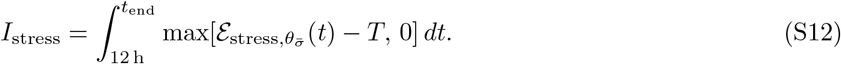

The calculation uses trapezoidal integration on the common 51-timepoint post-induction grid. Because elasticity is dimensionless, *I*_stress_ has units of hours; it is not an integral of fluorescence or a direct measure of accumulated cellular damage. Seven of the twelve genotypes have positive integrals and five have exact zeros under this rule. A zero denotes no excursion above the reference, not absence of a host interaction. The native AS20 and AS22 integrals are 2.5183 and 0.9443 h; AS20t has the smallest positive integral, 0.0090 h. Display labels round positive integrals to two decimals, with *<* 0.01 reserved for values that would otherwise round to zero; this is a display convention, not a statistical detection limit. Constant multiplicative reporter gain cancels from both the elasticity and its integral above a fixed threshold. Additive background, saturation, inferred rung spacing and biological context can still affect them. In particular, additive background can compress slopes at low signal, and a flat slope does not exclude a response shared across all RBSs.

#### Measured suppression score

The current potency panels in Fig. 5f,g use no Hill fit or inferred effective dose. For each sigma system, the dose-grid extraction subtracts a baseline estimated from the median reporter level in the no-vanillate condition and clips negative values to zero. Measurements at positive vanillate are pooled by their median across stress-promoter variants within each binder/RBS/salicylate combination, requiring at least two contributing variants. The suppression score is the number of these medians equal to zero among six RBSs and three salicylate levels (0, 25 and 100 *µ*M). Thus, “below detection” in these panels denotes the operational background-subtraction floor, not the independently estimated three-noise-standard-deviation limit used for the later kinetic experiment. The grid baseline also differs from the interpolated dark-window background used to display the reporter trajectories in panels c,d.

All 20 designs per target and the native and truncated controls appear in the potency–specificity plane. The displayed regions require cross-talk below 3 and at least five suppressed combinations for ECF20 or three for ECF22. These cutoffs summarize the selected set retrospectively; they do not establish a prospective selection procedure. Red rings identify the separate constitutive-mKate2Hyb burden exclusions. Previously explored Hill coefficients and half-maximal effective doses are not used in the current panels or their interpretation.

### S6 Optimal experiment design

Candidate designs comprise piecewise constant vanillate and salicylate levels and phase durations. Nominal models were the curated cascade fits obtained from earlier characterization of ECF20, ECF22 and SigW. The inspected joint-design driver uses equal system weights, pins *K*_*σ*_ and the observation background, and targets the shared parameters *β, η, γ*_*σ*_, 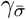, 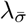. The expression ladders and sigma basal-expression parameters also enter the information calculation.

For log parameters *q* = log *ϑ*, define *s*_*ek*_ = *∂Ŷ* _*e*_(*t*_*k*_)*/∂q*. The implemented local information approximation is

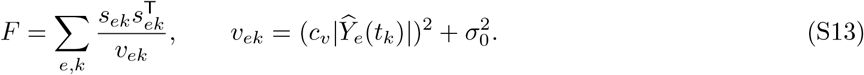

Sensitivities are computed by automatic differentiation ^61^. They are derivatives of fluorescence with respect to log parameters, not derivatives of log fluorescence. The proportional and additive noise terms are estimated from prior-model residuals. Equation S13 is the sensitivity-based information approximation used by the code; it does not include an additional information term from derivatives of the parameter-dependent variance.

For parameters of interest *I* and nuisance parameters *N*, the subset criterion uses the Schur complement ^53,54,62^

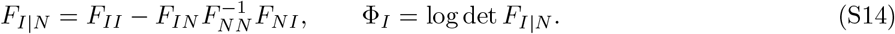

Pinned coordinates are removed first. A small diagonal regularization is added to the nuisance block for numerical evaluation, and near-singular designs are rejected. The selected 26-h schedule used ordinary *D*_*s*_-optimality, with the objective summed equally across nominal ECF20, ECF22 and SigW models. The selected shared kinetic parameters form the subset of interest; expression-ladder and basal sigma parameters are nuisance parameters. The search also includes a duration penalty. “Joint” here denotes the sum across systems, not the separate compound criterion implemented in the software. Both the generating job and the saved schedule metadata specify the ordinary *D*_*s*_ criterion.

The selected design record specifies a 26-hour budget, six variable phases, three-hour zero-inducer phases at both ends, vanillate in [0, 15] *µ*M, and salicylate in [0, 100] *µ*M. It records ten optimization starts and 500 iterations per start. The search implementation uses Nelder–Mead ^63^, bounded evaluation of candidate designs, and a fixed random seed of 42. The search driver evaluates a reduced RBS grid at 30-minute intervals; the final geometry export evaluates the full available grid at 15-minute intervals.

Concentrations and durations were rounded for experimental implementation:

| Phase | Vanillate ($\mu\text{M}$) | Salicylate ($\mu\text{M}$) | Duration (h) |
| --- | --- | --- | --- |
| 1 | 0 | 0 | 3.00 |
| 2 | 15 | 25.5 | 3.75 |
| 3 | 1.5 | 9 | 3.25 |
| 4 | 15 | 18.75 | 3.25 |
| 5 | 15 | 3.75 | 2.00 |
| 6 | 5.5 | 100 | 4.00 |
| 7 | 15 | 0 | 3.50 |
| 8 | 0 | 0 | 3.00 |

These durations sum to 25.75 h, within the nominal 26-h budget the design was optimized against. The final zero-input interval begins at 22.75 h and the last recorded sample is at 25.75 h. The table reports input intervals, not the number of sampled frames; actuation ends when the final zero-input interval begins, and the remaining 3 h is acquisition at zero input.

The saved geometry used in Fig. 6 was exported from the unrounded optimized schedule. Its ECF22 covariance submatrices give correlations of 0.940186 and −0.040857 for the conventional and optimized schedules, respectively, and a joint area ratio of 11.5752. These are local predictions under the nominal models and assumed noise. They are not empirical confidence regions. A common scalar change in observation noise rescales covariance without changing its correlation; changes in time dependence or signal-dependent weighting need not preserve either correlations or design rankings.

### S7 Dynamic inference from the designed experiment

The accepted result source for Fig. 6 is `260410 analysis/results/run final`. Sigma-only reference trajectories were fitted before the cognate binder trajectories. In the four-state binder model,

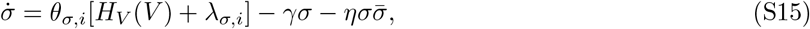

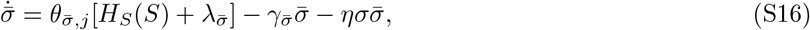

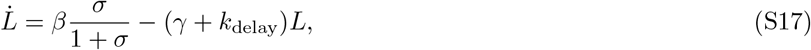

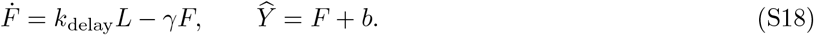

The reference model omits the antisigma state and sequestration terms. Reporter delay and inducer-response constants are supplied by the upstream reporter calibration. The reporter-specific removal rates from that calibration are replaced by the shared dilution rate specified below in both the sigma-only reference and cognate binder fits. The functions *H*_*V*_ and *H*_*S*_ have unit amplitude and exclude basal activity; the *λ* terms are additive. Their Hill coefficients are inherited from the reporter calibration, not fixed to one. Initial conditions are steady states under the initial input. The maximal-sequestration approximation supplies a warm start for the bimolecular binder fit.

The primary binder stage fits 15 parameters: twelve antisigma production activities, 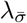,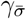, and *η*. The sigma ladder and its basal activities, reporter production scale, concentration gauge, and background are held fixed during this stage. The accepted run re-estimates the residual background before the cognate fit as the median of trajectory minima among the lowest four sigma and highest three antisigma indexed rungs, using unmasked observations. It falls back to the reference-fit background only if no such trajectory is available. Thus, this background is fixed during the cognate fit but is not generally the value from the noncognate reference fit. The accepted run uses locally background-subtracted fluorescence. For mVenus, observations through timepoint 26 are multiplied by the measured excitation-scale correction 1*/*2.5945 = 0.38544, not the nominal 0.4. Individual early mVenus frames with raw intensity above 45,000 are masked; camera saturation at 60,000 masks the trajectory from the first saturated frame onward. Observations before 1 h are also masked. The noncognate reference averages variant trajectories equally at each sigma rung; its saturation mask is based on the average raw signal, not on any single contributing variant. Low-signal cognate trajectories are retained in fitting rather than discarded because their residuals are small.

The weighted trajectory objective has the form of Eq. S4. For this run the residual-scale floor is pooled across the fitted grid. The scale of each trajectory remains its own standard deviation when this exceeds the pooled floor. Optimization uses L-BFGS with automatic differentiation in log-parameter coordinates ^61,64^; fixed coordinates are removed from the optimization vector. The saved run uses eight starts per architecture and a relative-loss convergence gate: two perturbed restarts per round for two rounds, with convergence declared when no restart improves the loss by more than 10^−3^ in relative terms. This is a loss-improvement criterion, not a gradient or parameter tolerance. All eleven reported binder architectures pass the recorded gate.

The fixed dilution rate is *γ* = log(2)*/*(28*/*60) = 1.4853154 h^−1^; the implemented antisigma lower bound is 0.999*γ* = 1.4838301 h^−1^. Seven binder estimates reach this bound. These are bound-limited removal estimates, and their sequestration estimates are conditional on that constraint. Neither the concentration normalization nor numerical convergence establishes absolute molecular concentrations or removes practical parameter correlations. For the correlation summary, a variant must have at least one retained observation above its channel-specific detection limit. The limit is three times the median across-trench standard deviation among variant–timepoint observations at the subtraction floor, requiring at least three trenches. The resulting limits are 37.1 A.U. for CFP and 128.7 A.U. for YFP. Eighty variants fail this reporting criterion; 1,453 variants from the 11 displayed architectures remain. Pearson correlation is calculated within each retained trajectory and summarizes temporal agreement, not agreement in absolute scale or offset.

The whiskers for *η* and 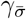 in Fig. 6f,g are conditional loss-slice ranges. Each parameter is varied on a grid while every other parameter is held at its fitted value. The endpoints delimit loss no greater than 1.01 times the minimum on that slice, with linear interpolation between grid points; an endpoint that does not cross this threshold is limited by the evaluated grid. The nuisance parameters are not reoptimized. These are neither profile-likelihood confidence intervals nor empirical 95% intervals, and they do not propagate uncertainty from the reference fit. They are distinct from the local design-stage Cramér–Rao regions ^65^.

### S8 Supplementary feedback theory: ideal integration and leakage

The preceding sections estimate open-loop component activities and kinetics. We now derive the closed-loop predictions used to interpret the feedback libraries, first for ideal integration and fast sequestration with leakage, and then for sequestration competing with independent removal. The subsequent measurement-analysis sections explain how these predictions are compared with experimental fluorescence.

The four-state dynamics are given in main-text Eq. 2. The states (*c*_1_, *c*_2_, *p, d*) denote free controller sigma, controller inhibitor, process sigma and disturbance inhibitor; the corresponding conventional AIF symbols are (*z*_1_, *z*_2_, *x, d*). At fixed induction, write *µ* = *µ*(*c*_V_) and Δ = *θ*_*d*_[*a*_SSal_(*c*_S_) + *λ*_*d*_]. The basal activity *λ*_*d*_ is specific to this circuit context, so the uninduced disturbance load need not vanish. Sequestration rates are *η*_*c*_, *η*_*d*_, and independent removal rates are 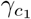, 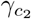, *γ*_*p*_, *γ*_*d*_.

#### Ideal integral balance

The ideal AIF controller is obtained when both controller species are removed only through mutual sequestration, such that 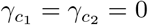. Subtracting the controller equations then gives

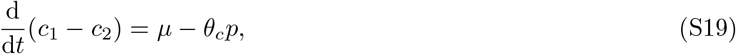

and any stable steady state must therefore satisfy the set point in Eq. 3, 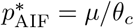. The process set point is consequently independent of the process-expression parameter *θ*_*p*_ and the disturbance-expression parameter *θ*_*d*_, while scaling inversely with the controller-expression parameter *θ*_*c*_.

#### Fast sequestration with controller leakage

Dilution and active turnover make the biological controller leaky. To expose the resulting departures from ideal adaptation, we consider the limit of fast sequestration on the controlled branch, where 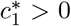 and the free concentrations of *c*_2_ and *d* are negligible. The steady-state balances then reduce to

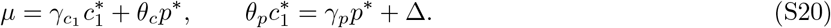

We define the dimensionless loop gain and disturbance load as

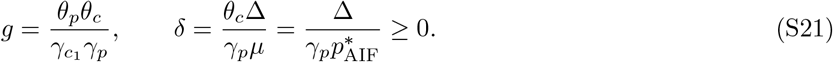

Thus, *g* compares closed-loop production with controller and process leakage, whereas *δ* compares disturbance production with process removal at the ideal AIF set point. Solving Eq. S20 gives

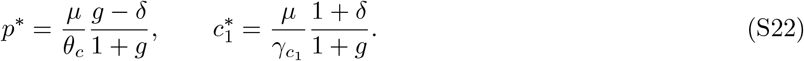

This branch exists for *g > δ*, equivalently

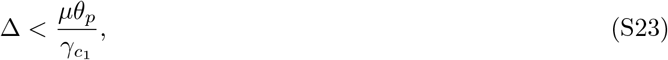

which defines the disturbance capacity of the leaky controller. As 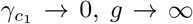, and the steady states converge to

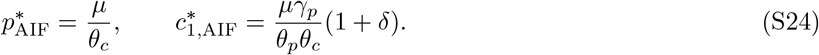

#### Elasticity predictions

Elasticities are defined in main-text Eq. 4; the controller index *c* refers to free *c*_1_. The derivatives below hold induction levels and other parameters fixed while varying the indicated expression activity. In particular, Δ scales with *θ*_*d*_, including its basal component.

Because both *g* and *δ* depend on the varied RBS activities, the six elasticities of Eq. S22 are

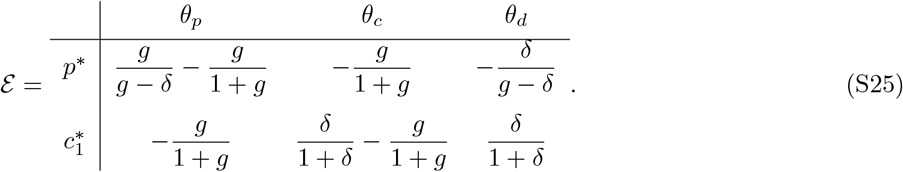

In the perfect-integrator limit *g* → ∞ at fixed *δ*, these become

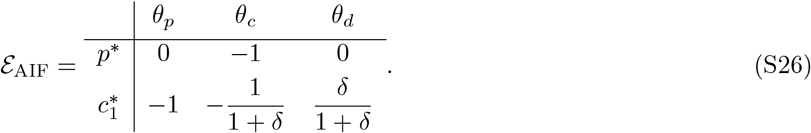

The ideal process is therefore insensitive to both its own expression parameter and disturbance strength, whereas the controller absorbs those variations through compensatory changes in 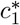. Importantly, basal disturbance expression gives *δ >* 0, so the ideal controller elasticities with respect to *θ*_*c*_ and *θ*_*d*_ need not equal −1 and 0, respectively.

### Scope of the reduced dynamics

On the controlled branch, the fast-sequestration approximation underlying Eq. S20 gives

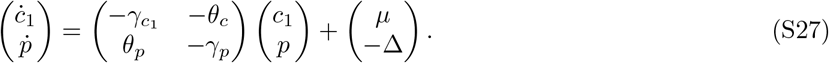

The disturbance enters only as an additive input. Changing its constant value therefore shifts the equilibrium without changing the Jacobian or its eigenvalues, provided the equilibrium remains on the same controlled branch and the approximation remains valid. A disturbance step can excite decaying oscillations, but cannot destabilize a previously stable equilibrium within this reduction. Disturbance-induced sustained oscillations consequently require dynamics beyond this description.

#### Equilibrium response beyond the fast-sequestration limit

The exclusion of equilibrium overadaptation is more general and does not require fast sequestration. Consider the full four-state model in Eq. 2, with positive kinetic parameters and fixed controller input *µ*. At equilibrium, the controller balances define free controller as a decreasing function of process abundance, 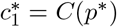. Specifically,

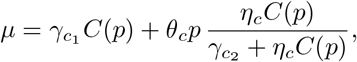

whose right-hand side increases with both *C* and *p*, implying *C*^′^(*p*) *<* 0. Eliminating *d*^*^ = Δ*/*(*γ*_*d*_ + *η*_*d*_*p*^*^) from the process balance gives

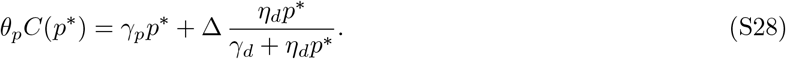

Implicit differentiation with all other inputs and parameters held fixed yields

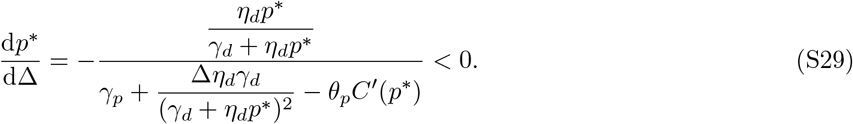

Thus, stronger disturbance lowers the equilibrium process output in the leaky model, irrespective of sequestration speed. In the ideal-integrator limit, 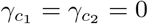, the controller balance instead fixes *p*^*^ = *µ/θ*_*c*_ wherever an equilibrium exists. Neither case permits an increase above the predisturbance equilibrium. This conclusion concerns equilibrium output; it does not exclude transient overshoot or establish a corresponding constraint on the means or medians of sustained oscillations.

### S9 Supplementary feedback theory: sequestration kinetics and controller responsiveness

To examine controller responsiveness without assuming complete sequestration, we retain both effective sequestration and independent removal in the controller balances of Eq. 2. Here, *η*_*c*_ is an effective sequestration parameter rather than a molecular association rate; its relation to reversible binding is developed at the end of this subsection. The steady-state balances are

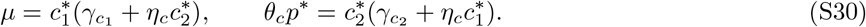

The first balance gives 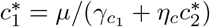. At a finite effective sequestration rate, residual free controller, and hence process production, can persist even when *c*_2_ *> c*_1_. Residual expression alone, however, does not determine how effectively feedback regulates the process.

The relevant quantities compare the competing removal routes of each controller species:

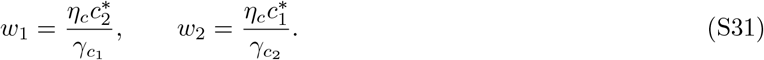

Thus, *w*_1_ measures sequestration relative to independent removal of free controller *c*_1_, whereas *w*_2_ measures sequestration relative to independent removal of its inhibitor *c*_2_. Large *w*_1_ means that most *c*_1_ molecules are sequestered rather than diluted or degraded, and similarly for *w*_2_ and *c*_2_.

At fixed controller input and kinetic parameters, the first balance constrains these ratios through *w*_2_(1 + *w*_1_) = Λ, where 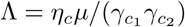. Treating process *p* as the input to the controller subsystem, implicit differentiation gives its local logarithmic response,

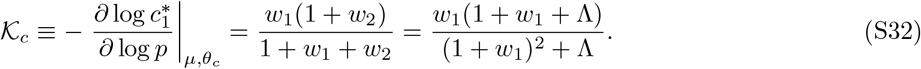

This is the controller response, not the full closed-loop gain. Its connection to the measured elasticities is 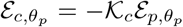 at equilibrium, with the other circuit parameters fixed: larger *K*_*c*_ means a larger controller response relative to the residual process response.

For Λ *>* 1, this expression has a unique maximum at an intermediate inhibitor loading:

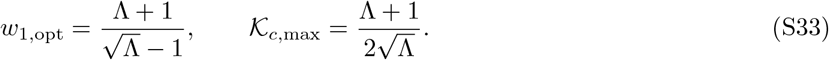

The controller response *K*_*c*_ increases below *w*_1,opt_ and decreases above it. The peak becomes large only when Λ ≫ 1; for Λ ≤ 1, the response instead rises monotonically toward unity and there is no finite-loading maximum. Increasing inhibitor production *θ*_*c*_*p*^*^ traverses this loading axis: the balances give *θ*_*c*_*p*^*^*/µ* = *w*_1_*/*(1 + *w*_1_) + *w*_1_*/*Λ, a strictly increasing function of *w*_1_. We therefore distinguish under- and over-actuated regimes with respect to the inhibitory *c*_2_ feedback branch, rather than the amount of free controller driving the process.

At low inhibitory loading (*w*_1_ ≪ 1), independent 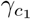-driven removal dominates the controller balance, 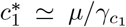, and K_*c*_ → 0. In this under-actuated regime, changes in *c*_2_ production have little effect on free controller. At excessive loading (*w*_1_ ≫ max{1, Λ} and *w*_2_ ≪ 1), sequestration consumes nearly all controller production, while independent 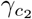-driven removal consumes most inhibitor production. Consequently, 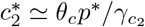 and 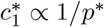, so *K*_*c*_ declines from its intermediate-loading maximum toward unity.

This regime is related to, but distinct from, integral windup as conventionally defined. Biomolecular windup arises when positivity or saturation prevents the process from realizing the actuation demanded by an integral controller, allowing the controller state to accumulate ^55^. Here, excess *c*_2_ suppresses free controller while its independent removal diverts inhibitor production away from the shared sequestration reaction. The inhibitor pool therefore becomes poorly coupled to the production-flux balance that supports near-ideal integration. We refer to this as a windup-like, over-actuated regime: it captures accumulation and reduced controller responsiveness, but does not require a separately saturating actuator or establish the prolonged transient recovery associated with classical windup.

When Λ is large, intermediate loading allows *w*_1_ and *w*_2_ to be simultaneously large. Sequestration then dominates removal of both species, approximately matching the two production fluxes, *µ* ≃ *θ*_*c*_*p*^*^. A disturbance that reduces process abundance and inhibitor production can move an initially over-actuated controller toward this more responsive regime. The theory therefore establishes an intermediate operating range for strong controller responsiveness and a mechanism for disturbance-dependent engagement. We do not assign individual library variants to these theoretical regimes or infer an experimental optimum from this calculation. The result concerns equilibrium responses and does not by itself establish dynamic stability.

#### Microscopic interpretation of effective sequestration

The controller-response calculation concerns the balance of effective sequestration and independent removal; it does not require molecular association itself to be slow or irreversible. To illustrate this distinction, consider a reversible controller–inhibitor complex *C* with association constant *k*_on_, dissociation rate *k*_off_, and first-order complex removal rate *γ*_*C*_, where removal of a complex removes both bound partners. At steady state, 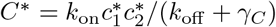. The net removal flux from either free species is then 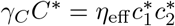, with *η*_eff_ = *k*_on_*γ*_*C*_*/*(*k*_off_ + *γ*_*C*_). When dissociation is much faster than complex removal, this becomes *η*_eff_ ≃ *γ*_*C*_*/K*_*D*_, where *K*_*D*_ = *k*_off_ */k*_on_. Rapid association and dissociation at finite affinity can therefore yield a finite effective sequestration rate and the same steady-state free-controller balances used above. This relation does not establish equivalence of the transient dynamics: an explicit complex pool can store and release bound proteins. We retain the reduced model for its tractable description of the measured responses, interpreting *η* as an effective parameter rather than assigning it uniquely to a microscopic binding mechanism.

### S10 Local fluorescence background and low-signal measurements

The closed-loop analyses in Figs. 4 and 7 use the reprocessed YFP and CFP background-subtracted measurement columns. The local estimator is calibrated using reporter-negative cells from the earlier reporter experiment. For each trench, segmentation masks identify contiguous rows at the upper and lower ends of the kymograph crop that contain no cell pixels at any sampled time. The larger of these two regions supplies the background pad. Let *q*_0.02,*c*_(*k, t*) and *Ī*_*c*_(*k, t*) be the second percentile and mean pixel intensity in that pad for channel *c*, trench *k*, and time *t*. The calibration is

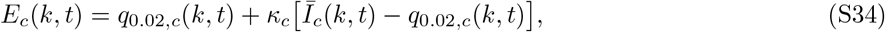

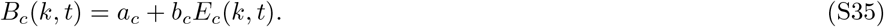

The stored coefficients (*κ*_*c*_, *a*_*c*_, *b*_*c*_) are (1, 62.8, 0.958) for CFP and (2, −52.9, 0.980) for YFP, with intensities expressed in camera counts. The calibration crop is 150 × 13 pixels, with a nominal trench length of 130 pixels and 20 pixels of axial padding. The estimator therefore depends on the crop geometry as well as the optical channel.

Calibration uses cells lacking the fluorescent-protein gene for the channel being measured. These cells provide a measurement of background over the segmented cell region, including contributions absent from an empty pad. The calibration analysis compares timepoint medians of pad estimates and over-cell intensity and fits the additive offset and slope by ordinary least squares. A separate comparison uses pads from adjacent bright trenches to predict the background in reporter-negative trenches. This comparison checks contamination associated with nearby fluorescence within the calibration experiment; it does not independently establish transfer of the calibration between experiments. Constitutive mKate2Hyb has no equivalent reporter-negative control and remains on its raw intensity scale.

#### Clipping and phase-level summaries

The reprocessed closed-loop tables retain raw intensity, estimated local background, and background-subtracted intensity. The latter is clipped at 1 count. A value at this floor therefore represents an upper bound on the untruncated difference. For Fig. 7, observations at or below this floor are counted before phase aggregation, and their counts and fractions accompany the phase-level estimates. The aggregation summarizes each trench in time and then takes the median across trenches. If a trench’s phase observations span multiple input storage blocks, the current extraction combines the block medians by an observation-count-weighted mean before the across-trench median. These counts describe clipping; they are not a fitted censored-data likelihood. The specific eligibility filters used for elasticities are given in Supplementary Section S11.

For Fig. 4, the extraction additionally defines a channel-specific low-signal threshold as 1.3 times the second percentile of background-subtracted measurements in the 0–3.5 h dark window. Phase levels below that threshold are flagged. This diagnostic threshold is distinct from the image-level subtraction and is applied to both channels before the full-ladder elasticity regression.

### S11 Experimental elasticities

For output *z* and expression activity *θ*_*j*_, following the elasticity convention of metabolic control analysis ^41–44^,

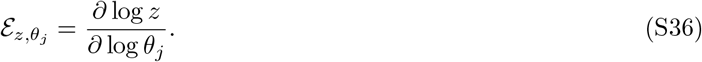

The experimental procedures estimate slopes from background-subtracted reporter fluorescence, with reporter inversion where specified below, against independently calibrated expression activities, holding the other RBS identities and architecture fixed. Both axes use the same logarithm base. Constant multiplicative units cancel, but additive background and reporter saturation do not. Each estimate is obtained within a fixed-background slice before any summary across slices. A full-ladder slope describes a finite sampled range; it need not equal the local derivative at every rung. The current main figures use these full-ladder regressions, not the moving-window local-elasticity estimates used in exploratory analyses.

For a first-order response *Y* = *Aσ/*(*K* + *σ*), with additive background already removed, inversion gives

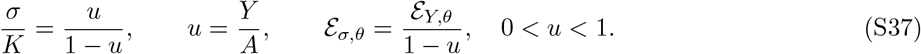

The last identity is local; a single correction factor need not transform a slope fitted over an entire ladder. Applying the inverse to the data requires a reporter ceiling on the same measurement scale.

#### Reporter inversion in Fig. 7 and Supplementary Fig. S7

All elasticity estimates in these figures invert the saturating reporter response before fitting log–log slopes. This correction is applied to both process and controller channels, using a separate fitted ceiling for each channel. It removes the response compression represented by the factor 1 − *u* in Eq. S37, including in the brighter open-loop reference. The inferred quantity *u/*(1 − *u*) is activity relative to the reporter’s fitted half-saturation scale; it does not establish an absolute molecular concentration. Agreement with complementarity is not used to choose the ceilings or to tune the estimator. The earlier analysis in Fig. 4c–e continues to use direct fluorescence slopes, without the reporter parameters inferred from the later designed experiment.

The steady-state reporter ceiling is

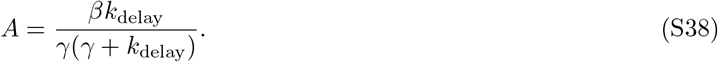

The preceding ECF22–AS22 fit supplies the process-channel production parameter (*β* = 67,416.1); the ECF20– AS20 fit supplies the controller-channel parameter (*β* = 18,395.7). Both ceilings use the shared removal rate imposed in those fits, *γ* = log(2)*/*(28*/*60) = 1.4853154 h^−1^, rather than the reporter-specific removal rates from the earlier calibration. The delay rate is reporter-specific: *k*_delay_ = 2.785 h^−1^ for the process reporter mVenus (`rcVenus` in the fit files), and *k*_delay_ = 6.933 h^−1^ for the controller reporter mJuniper. The process ceiling in the closed-loop experiment is *A*_*p*_ = *κA*_410_, where *κ* = 2.594458 is the measured 25%/10% excitation ratio read from `gain_estimate.csv` (`r_corrected`, 783 retained measurements). This is the same physical excitation ratio used in reverse to rescale the earlier experiment’s low-excitation segment. The controller channel was acquired at 100% excitation in both experiments and requires no corresponding gain rescaling. The resulting ceilings are approximately *A*_*p*_ = 76,800 and *A*_*c*_ = 10,200 fluorescence units. Using these ceilings across experiments assumes transfer of the fitted reporter response after the excitation correction; the gain measurement does not independently validate that biological transfer. Most fluorescence measurements lie well below their fitted ceilings, so reporter inversion primarily affects the brighter observations rather than typical signal levels.

#### Full-ladder regression in Fig. 4

Expression axes use the usable context-specific ladders from the sigma– antisigma calibration: ECF22 sigma for *θ*_*p*_, ECF20 antisigma for *θ*_*c*_, and ECF22 antisigma for *θ*_*d*_. The extracted levels are temporal medians of genotype-level, background-subtracted trajectories within schedule-derived windows. The extraction defaults to at least ten trenches and excludes the pTac arm. Within each fixed-background slice, ordinary least-squares slopes are fitted in log–log coordinates for both channels. Genotypes flagged as low signal in either channel are removed before regression; the flag is evaluated per genotype rather than per rung. A slice then requires positive outputs, at least six distinct activities, and at least 0.7 decades of input span. Reported summary elasticities are medians over qualifying slices. The low-signal thresholds remove no additional predisturbance rungs in this particular dataset; they nevertheless form part of the implemented filter. The process-, controller- and disturbance-expression sweeps retain 119, 143 and 118 slices, respectively. Their median process/controller slopes are (0.2934, −0.6491), (−0.6615, −0.5249) and (0.0125, −0.6406). Centering the plotted log levels within slices changes their intercepts, not these slopes; fitting a line through the pooled displayed points is not the estimator used for the reported elasticities.

#### Full-ladder estimates in Fig. 7c–g

The phase-level input table uses temporal medians within individual trenches, followed by medians across trenches for each genotype. The induced-phase windows are 6–12, 18–24, and 30–36 h. Activities are mapped from the independent designed-experiment fits, including binder-specific controller and disturbance ladders. The recovery window is 38–42 h, after salicylate withdrawal while vanillate remains present. The main analysis uses the ECF22-b16 disturbance architecture and requires at least five trenches, positive levels, and a calibrated process activity. Within each slice, process levels below 10% of the slice median are removed. Controller values below 40 fluorescence units are then removed rung by rung; they do not cause automatic rejection of the whole slice. At least six distinct process activities must remain after the signal filters and the ceiling exclusion described below. No missing value is imputed and no low-signal response is assigned zero elasticity. Rungs at or above 90% of either channel’s fitted ceiling are excluded before inversion, rather than clipped to a near-ceiling value: the inverse and its slope correction become unbounded as *u* → 1. This exclusion does not reduce the number of qualifying slices. For retained rungs, each channel is transformed to *Q* = *u/*(1 − *u*) using its own ceiling; a numerical lower guard sets *u* ≥ 10^−4^. Both slopes are estimated by Theil–Sen regression in log coordinates ^66,67^. For reporter-inferred activity *Q* in slice *s*, this is

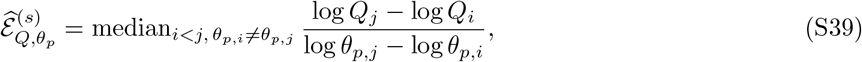

using the same retained rungs for the process and controller channels. Architecture summaries are medians of the process elasticity and absolute controller elasticity. Their 95% intervals are percentile intervals from 600 bootstrap resamples of paired slice estimates ^68^. This bootstrap conditions on the supplied expression calibration and treats slices as the resampling units; it does not propagate uncertainty in background, expression calibration, fitted reporter parameters or excitation ratio. Kernel-density contours, with bandwidth set by Scott’s rule ^69^, enclose the densest 50% of slice estimates and represent heterogeneity, not uncertainty in the median. There are 324, 388, 388 and 316 qualifying slices in the four successive windows. Complementarity is summarized as the median of the per-slice sums 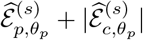, not a sum of separately pooled medians. The four successive values are 1.0961, 1.1023, 1.0651 and 1.1146. Eligibility is reassessed in each phase, so these summaries do not measure paired changes in an identical set of slices. They also include oscillatory trajectories, whose phase medians need not represent equilibria. Supplementary Fig. S7 applies the same procedure separately to each disturbance partner; binder medians require at least four slices and density contours at least eight.

#### Process-ladder visualization and open-loop reference in Fig. 7a

This panel uses the same predisturbance closed-loop slices, signal filters and channel-specific inversions as the paired analysis above. The process levels are inverted once, before both slope estimation and visualization; no additional correction is applied to the plotted closed-loop ladders. The analysis reports the median within-slice Theil–Sen slope of reporter-inferred process activity against the calibrated process-expression parameter. For visualization, each slice is normalized to its weakest retained process rung and medians are taken at each available activity. Slice membership can therefore differ along a displayed curve; the legend slope is not a regression through these aggregated curve points. The controller floor preferentially truncates the high-process-activity end of slices with strong controller compensation. Consequently, these slopes describe the retained activity ranges and should not be read as uniform-coverage estimates over the entire designed ladder.

The open-loop reference uses ECF22 variants carrying noncognate ECF20 binders in the preceding kinetic experiment. It summarizes the last four recorded timepoints in the 6–12 h interval, retains fixed binder/RBS slices with at least five process rungs, and uses the independently fitted sigma activity ladder and *A*_410_. It is not a matched feedback-deletion control from the closed-loop experiment. The median corrected slope is 1.05. Repeating the closed-loop transformation at *κ* = 1.5 and 6 preserves the order ECF20-b19, AS20t, AS20, ECF20-b6, followed by ECF20-b18 and ECF20-b11, whose relative order changes. The two latter binders are therefore not assigned a resolved ordering.

#### Kinetic association

Figure 7g correlates the logarithm of the independently fitted ECF20 binder ratio 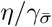 with the median absolute predisturbance controller slope for each binder. The Pearson calculation uses six binder medians, not the individual slice points; the Spearman calculation uses their ranks. Reported *P* values are two-sided. No uncertainty in the kinetic point estimates is propagated. These correlations are associations across six architectures, not fitted predictions of individual circuits.

### S12 Lineage autocorrelation and oscillation prevalence

**Autocorrelation curves in Fig. 4**. Dynamics are calculated within individual trench trajectories before averaging across trenches. For the illustrated genotype, the windows are 5.5–12.25 h before disturbance, 13–24.75 h at 25 *µ*M salicylate, 25.5–36.75 h at 50 *µ*M, and 37.5–44.75 h during recovery. Trajectories require at least 20 observations in a window. Background-subtracted intensity is clipped at 1 count, transformed to log_10_, and detrended by subtracting a fitted straight line in observation index.

At each positive lag, the autocorrelation is the Pearson correlation between the overlapping leading and trailing trajectory segments. The requested lag range extends to 6 h, with a shorter range for trajectories lacking enough observations; correlations require at least four overlapping pairs. The plotted curve is the mean of available lineage correlations. Shading is one standard error, calculated as the population standard deviation of the lineage correlations divided by the square root of the number of lineage curves in that phase. This display does not provide simultaneous confidence bands across lags.

A descriptive period is calculated only when the mean autocorrelation has at least three sign changes over the displayed range. The period estimate is twice the lag of its first strict local minimum. This is a descriptive criterion for repeated ringing. Residual curvature during recovery can produce a broad trough and a long-lag rebound after linear detrending, so a trough alone is not used to assign a period.

#### Per-trench prevalence in Fig. 7

The prevalence calculation uses raw YFP trajectories sampled at 15-minute intervals. It excludes the first 2.5 h of each induced phase, giving the half-open windows [6.5, 12), [14.5, 24), and [26.5, 36) h. A trench contributes when its window contains at least ten observations and all of them are finite, so a trench with any missing frame in the window is dropped entirely; a variant requires at least three contributing trenches. Each trajectory is linearly detrended in observation index. For a detrended trajectory *x*_1_, …, *x*_*n*_, the normalized autocorrelation is

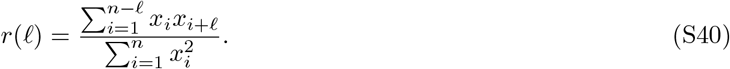

The denominator is common to all lags; it is not renormalized for the decreasing number of pairs. Among local autocorrelation maxima at lags from 1.3 to 6 h, the highest peak is selected. A trench is classified as oscillatory when that peak exceeds 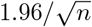 traces without an in-band peak are classified as nonoscillatory. The per-variant prevalence is the fraction of contributing trenches that meet this rule.

This threshold uses a pointwise white-noise reference. It is an operational classifier and does not account for colored noise or selection among multiple lags. The implementation treats consecutive retained observations as 15 minutes apart and does not interpolate missing frames. The prevalence used in the figure is computed from this autocorrelation classifier; the separate spectral detector in the same analysis does not gate the prevalence estimate.

#### Operational stability boundaries

For the ECF20-b19/ECF22-b16 heatmap, each process/controller RBS cell is the median per-variant oscillation fraction across disturbance-expression rungs. For the controller-expression profile, the median is instead taken across all scored variants at each controller rung. The profile is ordered by calibrated controller activity, which need not follow the nominal order of the RBS labels. A rung is classified as oscillatory when its median prevalence is at least 0.30. The boundary is the next sampled activity above the last such rung, ensuring that every higher observed rung is below the cutoff. If all rungs are below 0.30, no dividing boundary is drawn; if the largest sampled activity remains above the cutoff, no stable upper region is identified within the measured range.

Boundaries are determined separately for each phase in the ECF20-b19/ECF22-b16 comparison. For the comparison across controller binders, each binder uses its own calibrated activity ladder and its own boundary at 25 *µ*M salicylate, with the ECF22-b16 disturbance binder held fixed. These boundaries are empirical summaries of oscillation prevalence over the sampled library, rather than fitted bifurcation points of the dynamical model. In Supplementary Fig. S8, the analysis is repeated separately for each controller–disturbance pair at 25 *µ*M salicylate. Heatmap cells require at least three scored variants. Pair-level summaries require at least 100 variants; boundary profiles require at least six controller rungs with at least ten scored variants per rung. Missing or insufficiently sampled combinations are shown explicitly. Both heatmap axes are ordered by their calibrated activities; the main heatmap retains the nominal RBS-rung order. Pooling across disturbance-expression rungs is used only for these prevalence summaries and is not an assumption of invariance to disturbance expression.

### S13 Software

The Python analyses use NumPy ^70^ and SciPy ^58^; optimization and stiff integration are `scipy.optimize.least squares` and `scipy.integrate.solve_ivp` from the latter. The trajectory fits for the designed experiment are implemented in Julia ^71^, with sensitivities from ForwardDiff.jl ^61^ and quasi-Newton optimization from Optim.jl ^64^.

Code and analysis scripts are still being prepared for publication at the time of this preprint and will be made publicly available at a later date.

## Supplementary notation

**Table S1.**
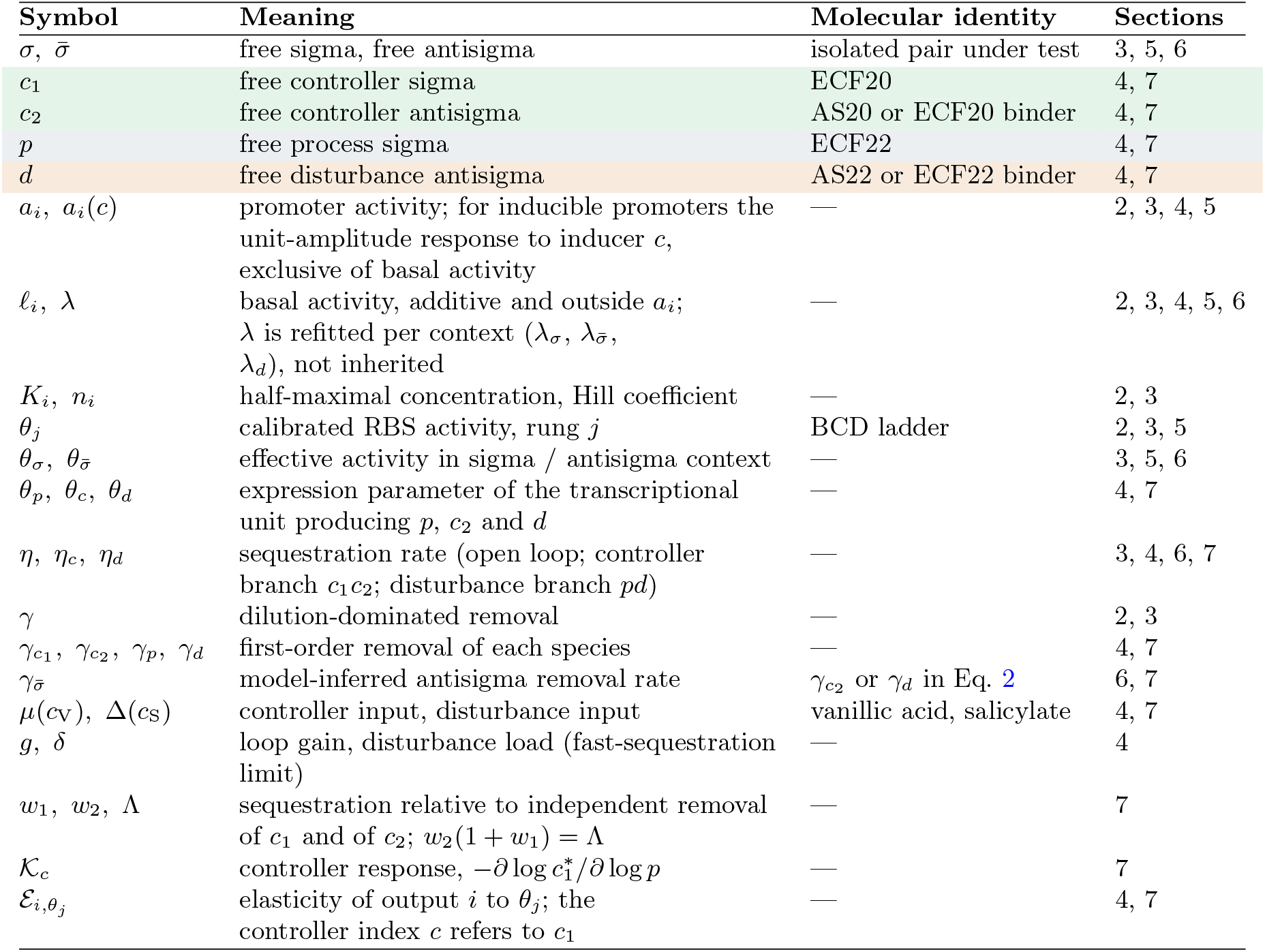
Notation. Free concentrations are written as 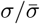 in the open-loop characterization sections and by circuit role in the closed-loop sections. For stress elasticities in Section 5, 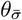 denotes the reference RBS activity *θ*_*j*_ driving binder expression, rather than a protein-specific activity inferred from kinetic fits.

| Symbol | Meaning | Molecular identity | Sections |
| --- | --- | --- | --- |
| $\sigma, \bar{\sigma}$ | free sigma, free antisigma | isolated pair under test | 3, 5, 6 |
| $c_1$ | free controller sigma | ECF20 | 4, 7 |
| $c_2$ | free controller antisigma | AS20 or ECF20 binder | 4, 7 |
| $p$ | free process sigma | ECF22 | 4, 7 |
| $d$ | free disturbance antisigma | AS22 or ECF22 binder | 4, 7 |
| $a_i, a_i(c)$ | promoter activity; for inducible promoters the unit-amplitude response to inducer $c$, exclusive of basal activity | — | 2, 3, 4, 5 |
| $\ell_i, \lambda$ | basal activity, additive and outside $a_i$; $\lambda$ is refitted per context ($\lambda_\sigma, \lambda_{\bar{\sigma}}, \lambda_d$), not inherited | — | 2, 3, 4, 5, 6 |
| $K_i, n_i$ | half-maximal concentration, Hill coefficient | — | 2, 3 |
| $\theta_j$ | calibrated RBS activity, rung $j$ | BCD ladder | 2, 3, 5 |
| $\theta_\sigma, \theta_{\bar{\sigma}}$ | effective activity in sigma / antisigma context | — | 3, 5, 6 |
| $\theta_p, \theta_c, \theta_d$ | expression parameter of the transcriptional unit producing $p, c_2$ and $d$ | — | 4, 7 |
| $\eta, \eta_c, \eta_d$ | sequestration rate (open loop; controller branch $c_1c_2$; disturbance branch $pd$) | — | 3, 4, 6, 7 |
| $\gamma$ | dilution-dominated removal | — | 2, 3 |
| $\gamma_{c_1}, \gamma_{c_2}, \gamma_p, \gamma_d$ | first-order removal of each species | — | 4, 7 |
| $\gamma_{\bar{\sigma}}$ | model-inferred antisigma removal rate | $\gamma_{c_2}$ or $\gamma_d$ in Eq. 2 | 6, 7 |
| $\mu(c_V), \Delta(c_S)$ | controller input, disturbance input | vanillic acid, salicylate | 4, 7 |
| $g, \delta$ | loop gain, disturbance load (fast-sequestration limit) | — | 4 |
| $w_1, w_2, \Lambda$ | sequestration relative to independent removal of $c_1$ and of $c_2$; $w_2(1 + w_1) = \Lambda$ | — | 7 |
| $\mathcal{K}_c$ | controller response, $-\partial \log c_1^* / \partial \log p$ | — | 7 |
| $\mathcal{E}_{i,\theta_j}$ | elasticity of output $i$ to $\theta_j$; the controller index $c$ refers to $c_1$ | — | 4, 7 |

## Supplementary figures

**Figure S1.**
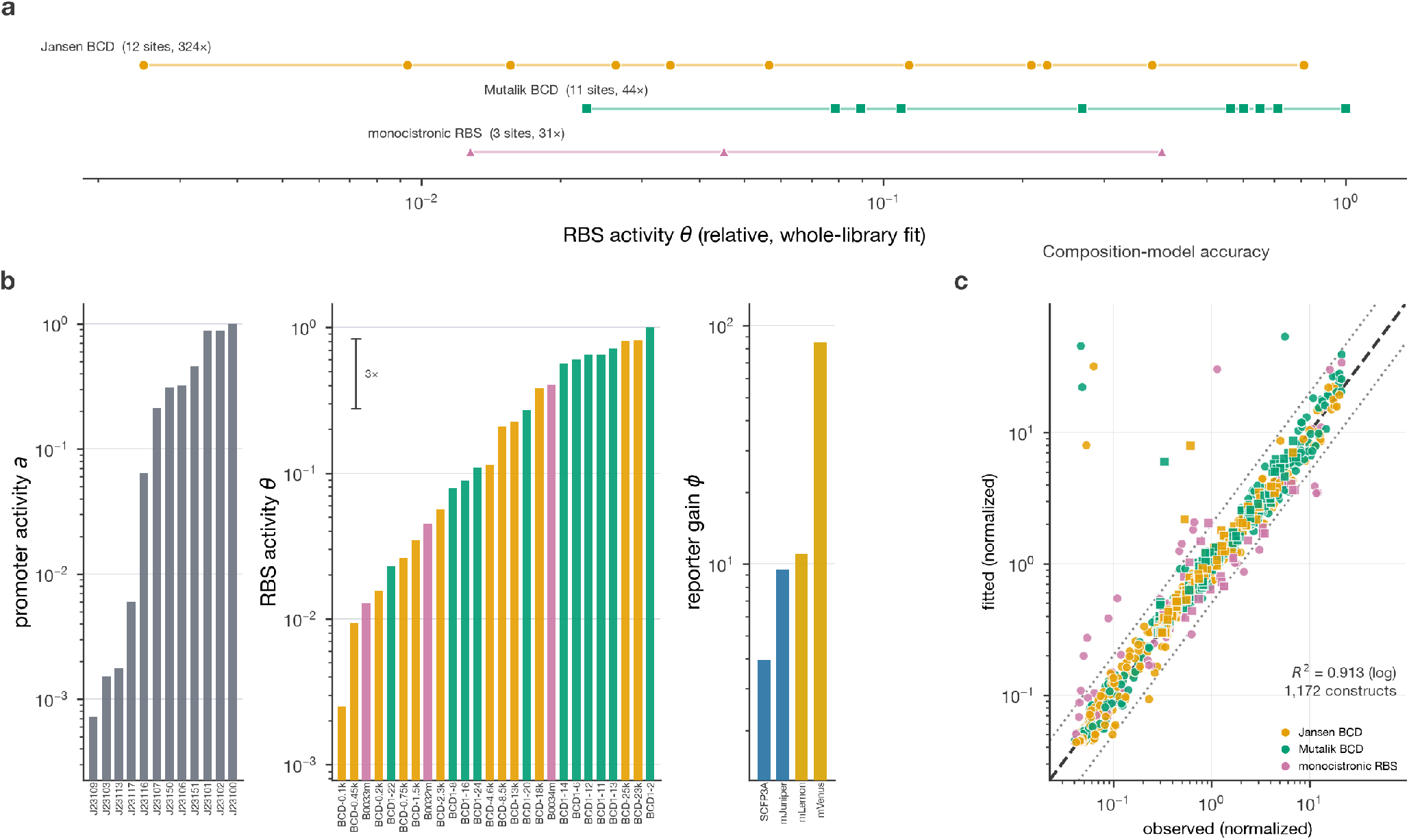
Calibration across three RBS design families. **a**, Activities of 26 RBSs from a joint whole-library fit, grouped by family; lines span each family’s range. **b**, Fitted activities of 12 constitutive promoters and 26 RBSs, and gains of four fluorescent reporters. **c**, Observed versus fitted normalized fluorescence for 1,172 constructs (*R*^2^ = 0.913, log space). Colors denote RBS families; circles/squares denote YFP/CFP-channel reporters. Dashed and dotted lines show identity and twofold deviations. The whole-library fit re-estimates shared activities and is distinct from the BCD-only fit in Fig. 2.

**Figure S2.**
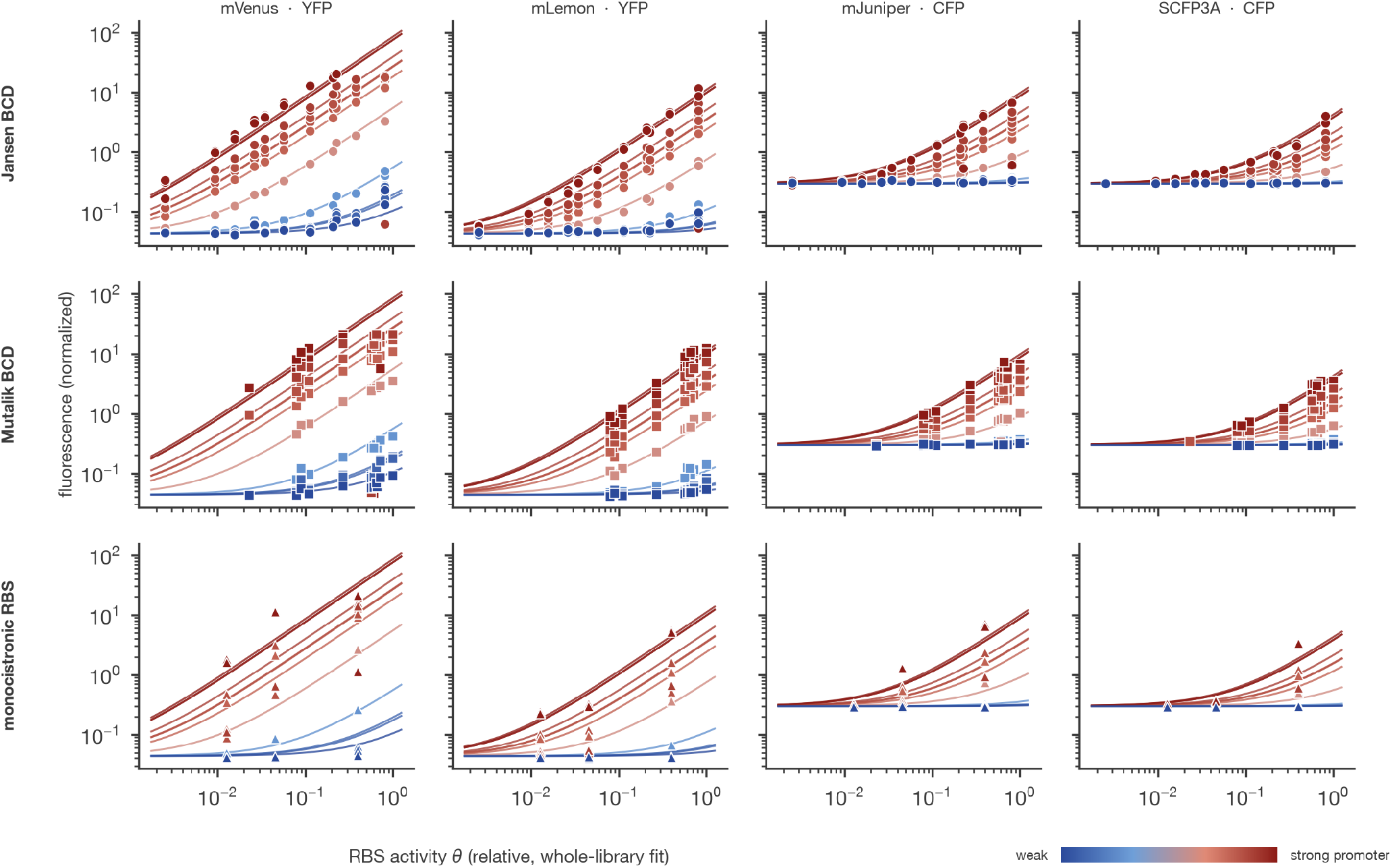
Reporter-context dependence across the three RBS families. Rows denote RBS families and columns denote fluorescent reporters. Points are construct measurements normalized to constitutive mKate2Hyb in the same cell and frame; lines are predictions of the joint composition model used in Supplementary Fig. S1, colored by promoter activity. All panels share the same fitted RBS axis, with no reporter-specific RBS refitting. Systematic departures reveal context dependence beyond the separable model.

**Figure S3.**
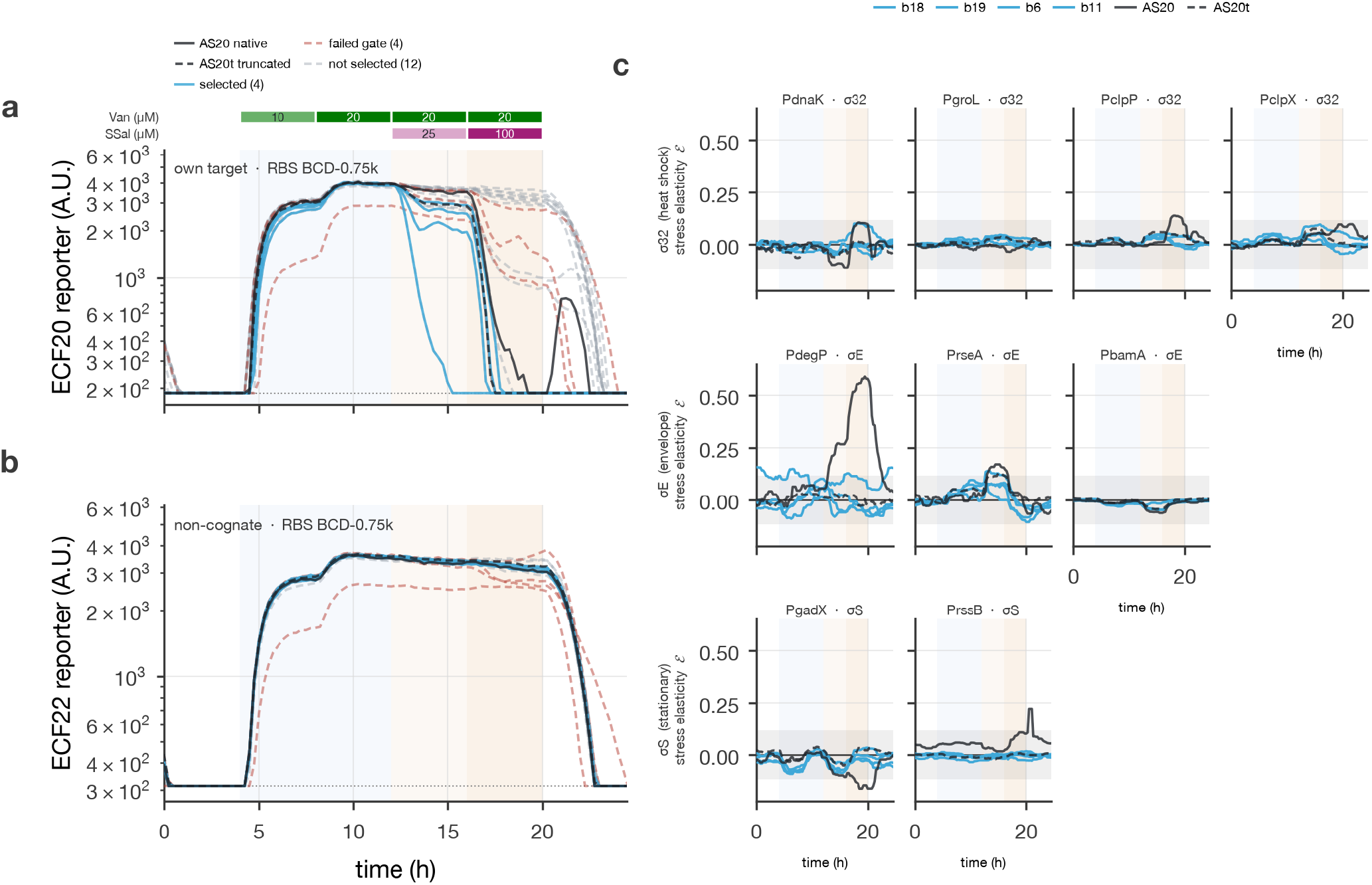
Cognate suppression, specificity and stress responses of ECF20 binders. **a,b**, Variant-median cognate ECF20 and noncognate ECF22 reporter trajectories at binder RBS BCD-0.75k, extending Fig. 5c,d to ECF20 binders. Blue, selected designs; black solid/dashed, native/truncated AS20; red dashed, failed specificity gate; gray, other designs. Dotted lines indicate display floors. **c**, Stress elasticities across all six RBSs, using the shared constitutive-expression reference activities from Fig. 2 and grouped by associated regulon. Gray bands mark *±*0.1175, the same reference threshold used in Fig. 5e and Supplementary Fig. S4. All nine promoters are retained, including weakly expressing *P*_bamA_. All four selected binders share a color to emphasize their comparison with native AS20; individual identities are not distinguished by color.

**Figure S4.**
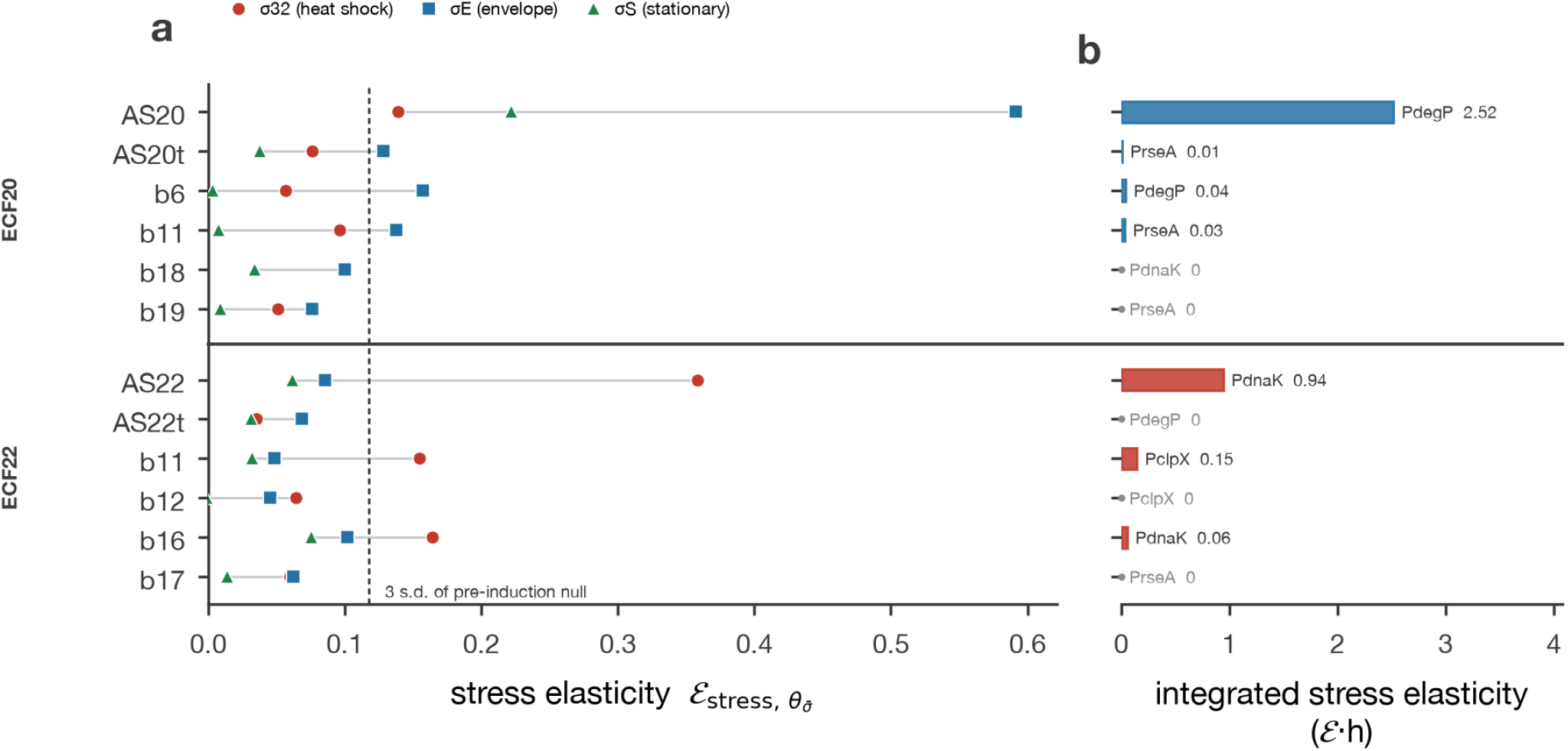
Native antisigma factors engage distinct stress responses. **a**, Maximum signed stress elasticity within each regulon over the recorded time course. All genotypes use the same constitutive-expression RBS reference activities from Fig. 2. The dashed line is three times the pooled 5–11 h reference standard deviation (threshold 0.1175). **b**, Positive elasticity above this threshold integrated from 12 h onward on the promoter carrying each binder’s largest post-induction peak. Color denotes regulon; labels name the promoter and report the integral in hours. Five gray zeros indicate no above-threshold post-induction excursion, not missing data. Native AS20 and AS22 integrals are 2.52 and 0.94 h; AS20t has a small positive integral (0.009 h). Two of four selected binders per target exceed the threshold, with smaller integrated responses than their native controls.

**Figure S5.**
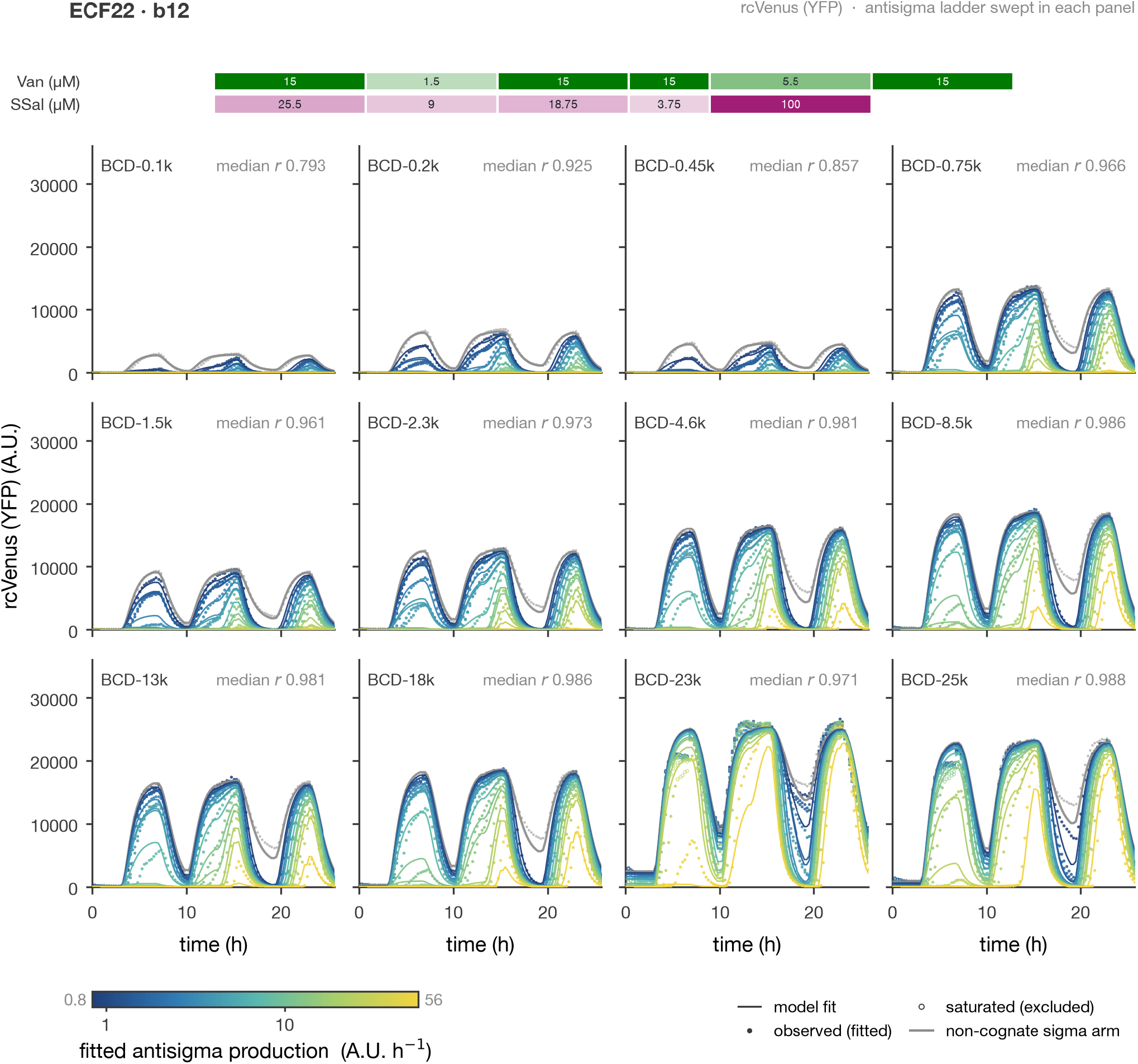
ECF22-b12 dynamic fits across both expression ladders. Each panel fixes one ECF22 RBS and varies the 12-member ECF22-b12 RBS ladder, extending the BCD-8.5k slice in Fig. 6e. Filled points are fitted observations; open points were excluded from the loss. Colored lines are joint-model fits, with color indicating fitted antisigma production activity. Gray curves show the noncognate reference arm, not an empty-vector control. Each annotated correlation is the median across that panel’s antisigma trajectories, using retained observations. All panels share a zero-based fluorescence scale. Fits for all 11 architectures are provided in Supplementary Data 1.

**Figure S6.**
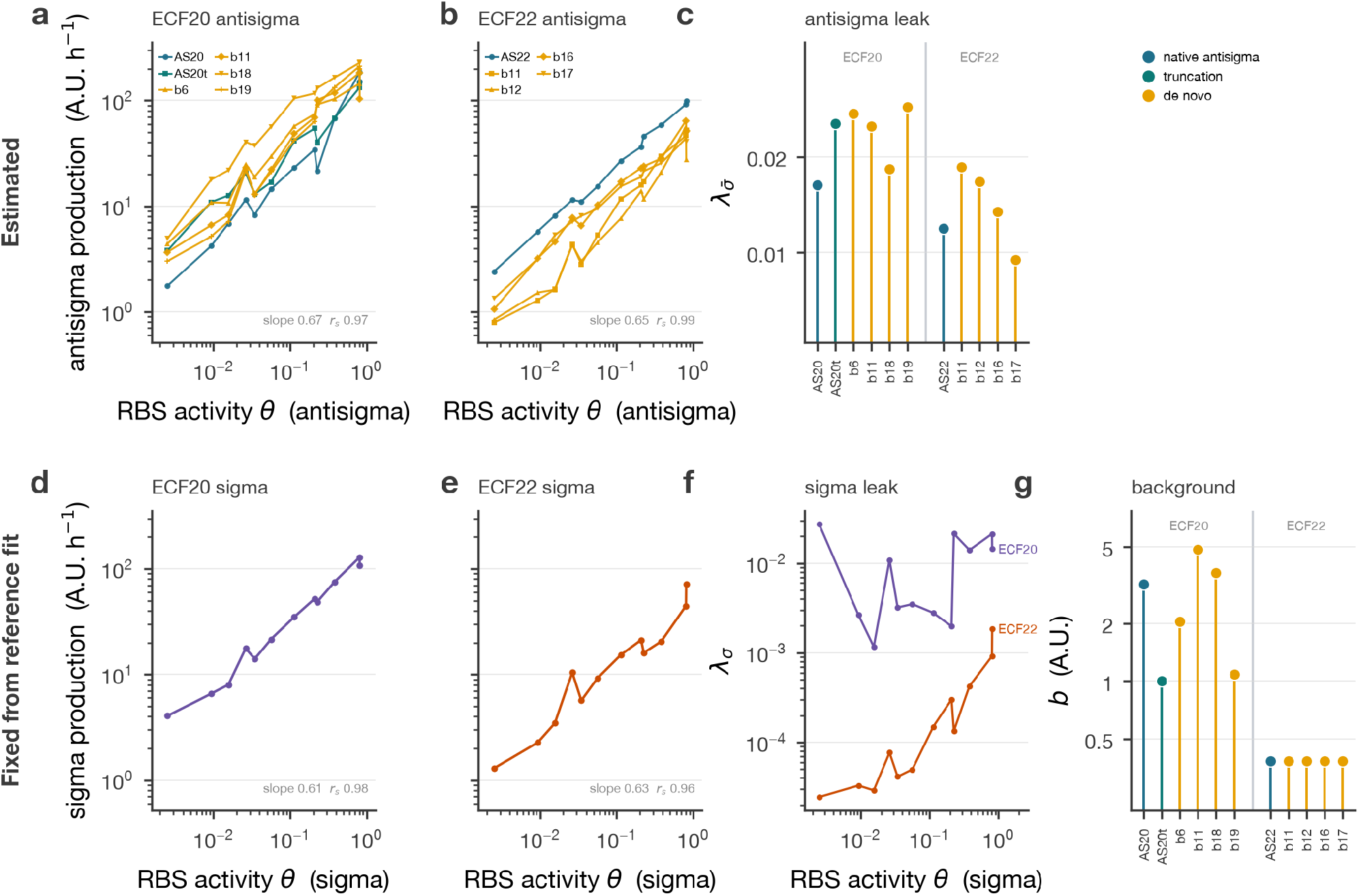
Expression and basal-activity parameters from the designed experiment. **a,b**, Fitted antisigma production activities for ECF20 and ECF22 binders versus the constitutive RBS calibration. **c**, Fitted antisigma basal activities. **d–f**, Sigma production ladders and RBS-dependent basal activities, obtained from noncognate reference fits and fixed during cognate fitting. **g**, Residual backgrounds fixed during cognate fitting after estimation from strongly sequestered trajectories. These need not equal the noncognate reference-fit backgrounds. For all five ECF22 architectures, and for AS20t, the estimate coincides with the one-count subtraction floor after excitation rescaling and should be read as an upper bound rather than a resolved value. Lines connect RBSs, not independent kinetic experiments. Activities are in system-specific model units and do not establish absolute molecular concentrations.

**Figure S7.**
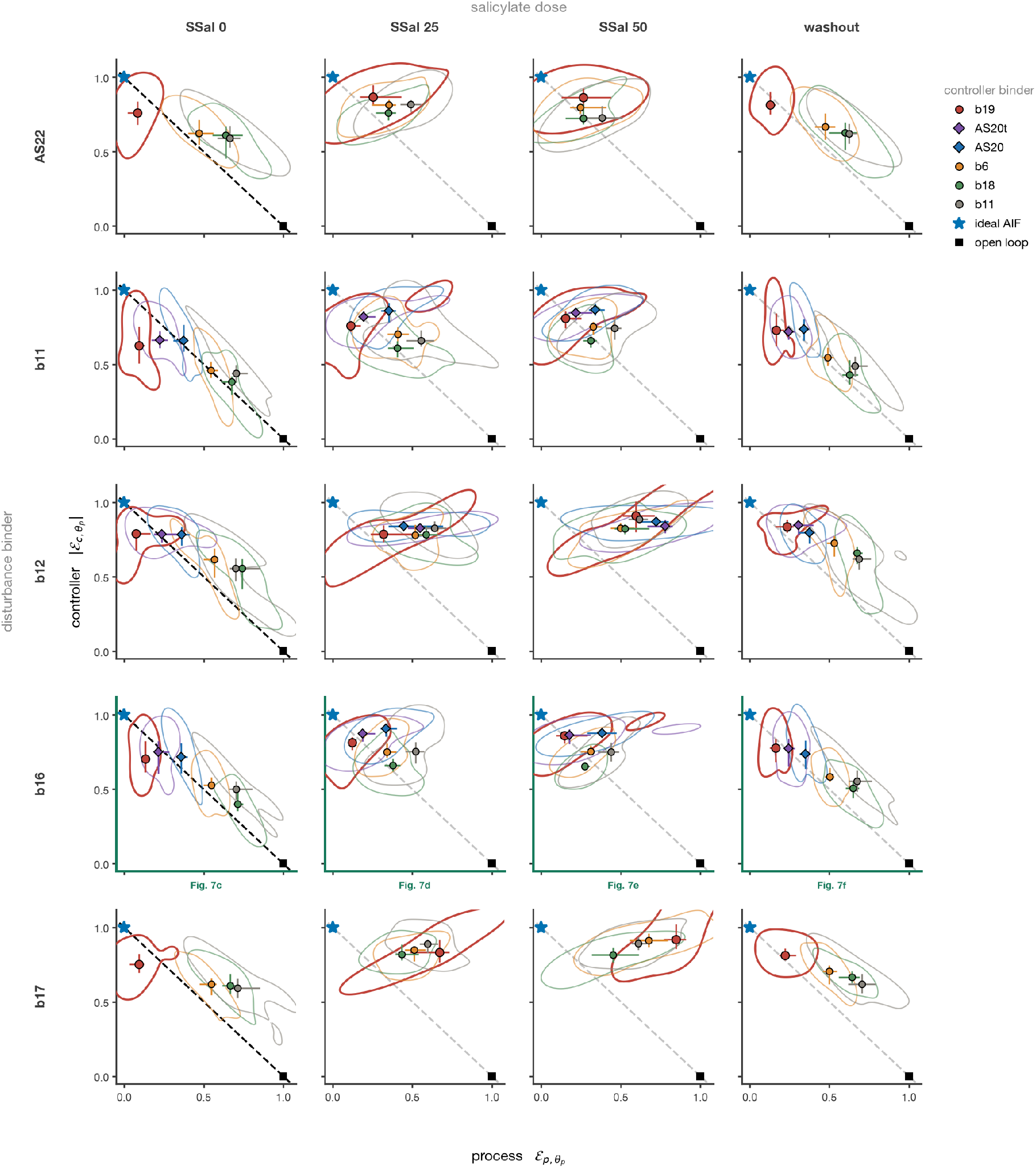
Process–controller sensitivity across disturbance architectures and conditions. Columns denote predisturbance, 25 and 50 *µ*M salicylate, and salicylate withdrawal; rows denote disturbance binders. Within each cell, colors identify controller binders. Symbols are median paired activity elasticities after channel-specific reporter inversion, whiskers are 95% slice-bootstrap intervals, and contours enclose the densest 50% of slice estimates. The estimator and signal-floor and near-ceiling exclusions match Fig. 7c–f, reproduced in the ECF22-b16 row. Medians require at least four qualifying slices and contours at least eight; missing combinations reflect insufficient coverage. The diagonal connects ideal AIF and open loop as a zero-disturbance reference, not an exact constraint under finite disturbance. Eligibility is reassessed in each condition.

**Figure S8.**
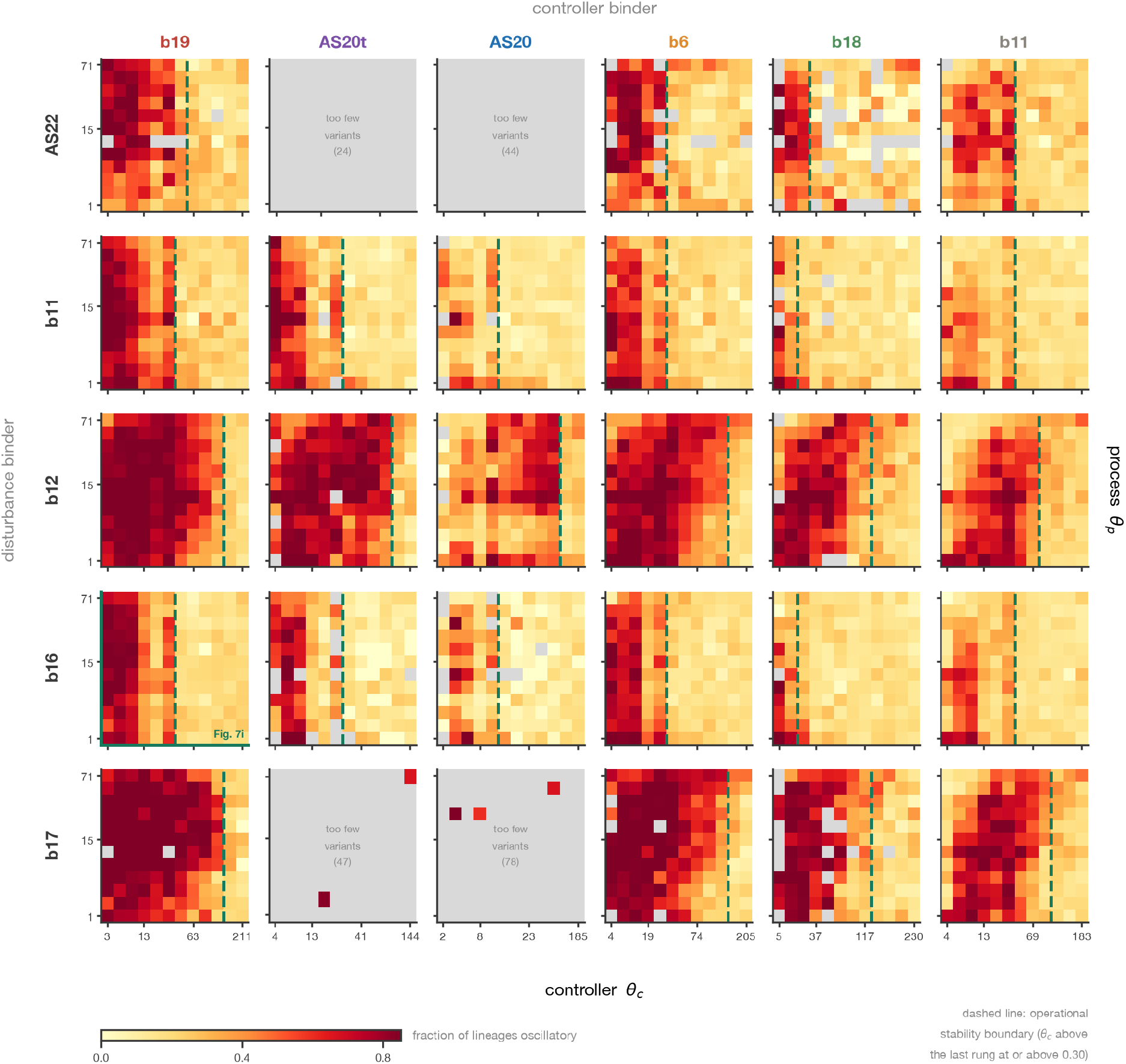
Oscillation prevalence depends on both controller and disturbance architecture. Per-lineage oscillation prevalence at 25 *µ*M salicylate across all six controller binders (columns) and five disturbance binders (rows). Each cell is the median per-variant prevalence across disturbance-expression rungs at fixed process and controller RBSs; at least three scored variants are required. Axes are ordered by calibrated activities. Dashed lines mark operational boundaries above the last controller rung with median prevalence at least 0.30, not fitted bifurcations. Gray cells or annotated panels indicate insufficient sampling. The ECF20-b19/ECF22-b16 panel corresponds to Fig. 7i, with its axes reordered by calibrated activity. The ECF22-b16 disturbance series is comparatively quiescent and should not be taken as representative of all partners.

**Figure S9.**
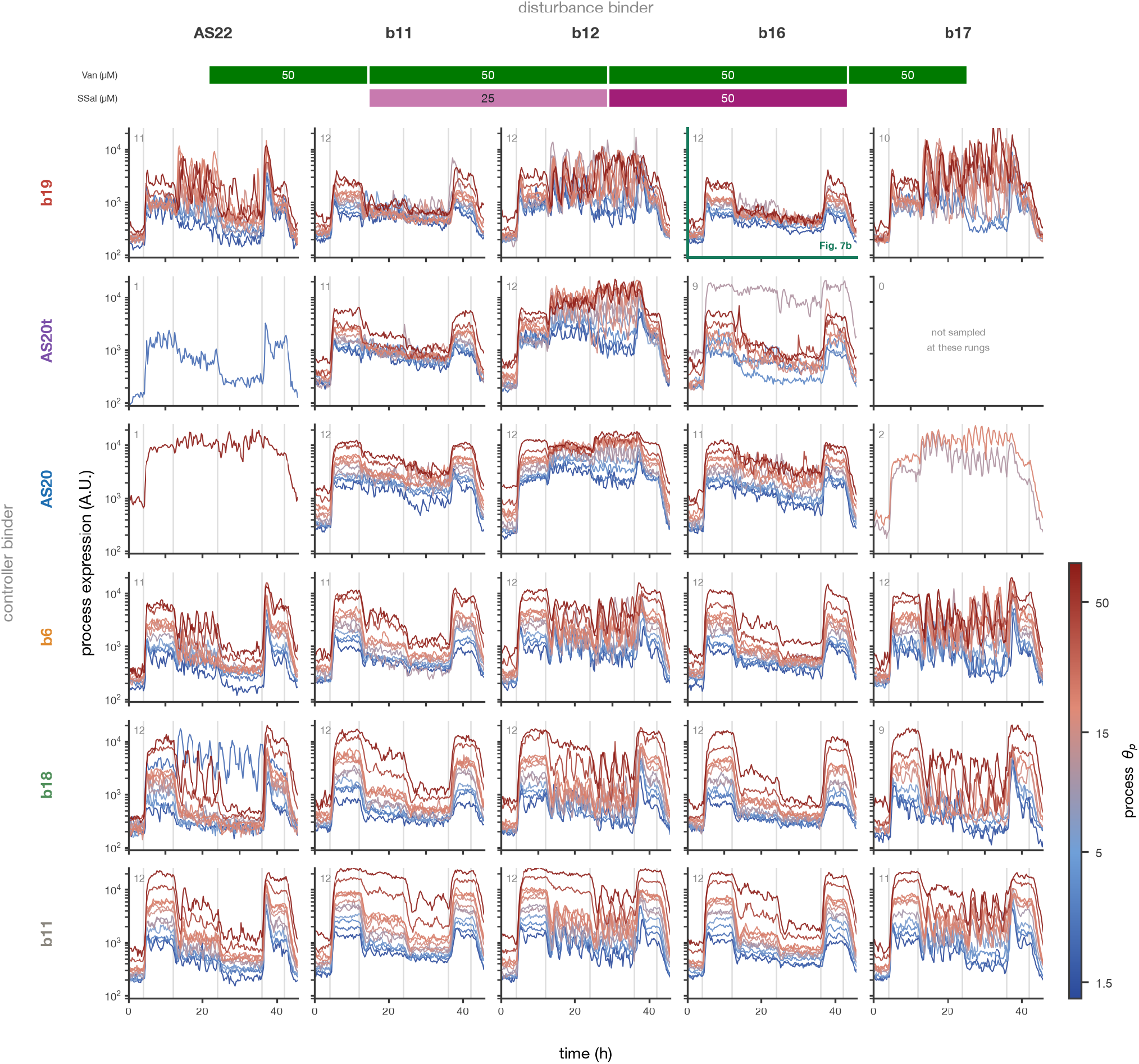
Matched process-expression sweeps across all binder pairs. Variant-median process-reporter trajectories for each controller (rows) and disturbance binder (columns), holding controller and disturbance RBSs at BCD-4.6k and BCD-2.3k while varying the process RBS. Colors denote calibrated process activity on a common scale. Numbers give the displayed rung count; each variant requires at least three trenches. The green-framed ECF20-b19/ECF22-b16 panel reproduces the process traces in Fig. 7b. All panels share the input schedule and fluorescence limits. Missing traces indicate insufficient sampling, not demonstrated absence of a phenotype. Median traces can obscure asynchronous lineage oscillations; lineage-resolved prevalence is shown in Supplementary Fig. S8.

